# Casp1 and Ripk3 are required for homeostatic insulin secretion in mice

**DOI:** 10.64898/2026.09.01.748578

**Authors:** Madison D. Girouard, Conor O’Dwyer, Cassandra A.A. Locatelli, Myriam P. Hoyeck, Evgenia Fadzeyeva, Tyler K.T. Smith, Rui Yan Gao, Tanvi Ahluwalia, Sophia M. Perrakis, Andrew C. Clément, Antonio A. Hanson, Peyman Ghorbani, Lili Grieco-St-Pierre, Jianfan I. Nie, Mariam Hakoum, Julia R.C. Nunes, Aaron Reyes, Andrew R. Pepper, Subash Sad, Jennifer E. Bruin, Erin E. Mulvihill, Morgan D. Fullerton

## Abstract

**Objectives:** Cell death and inflammatory pathways play important roles in adaptations to nutrient overload and metabolic dysfunction. This study investigates the metabolic consequences that arise from the dual disruption of both caspase 1 (Casp1) and receptor interacting protein kinase 3 (Ripk3) in mice fed a control or obesity-inducing diet.

**Methods:** Male and female wild-type (WT), Casp1/11 knockout (KO), Ripk3 KO and Casp1/11/Ripk3 double knockout (DKO) mice were fed a matched low-fat or a 60% kcal high fat diet, followed by metabolic phenotyping. Islets were isolated from WT and DKO mice for measures of dynamic glucose-stimulated insulin and somatostatin (Sst) secretion. Islet architecture and cellular composition were assessed in WT and DKO mice by immunofluorescent staining of intact pancreatic sections. Pharmacological inhibition of Casp1 (Ac-YVAD-cmk) and Ripk3 (GSK’872) was performed in WT and DKO mice using isolated islets and *in vivo* administration. Exogenous hormones were administered prior to glucose injection to test in vivo responses.

**Results:** High-fat feeding resulted in increased adiposity in male, but not female mice, with single or double deletion of Casp1/11 and Ripk3. These mice also exhibited markers of impaired glucose tolerance and insulin sensitivity. Interestingly, when both Casp1 and Ripk3 were deleted or inhibited in mice fed a low-fat diet, mice experienced reductions in glucose excursion following administration of glucose due to increased plasma insulin levels. This increase in insulin secretion was recapitulated in isolated islets *ex vivo* and was independent of changes in the proportions of α-, β-, and δ-cells within the islet. There were significant reductions in the percentage of urocortin-3 (Ucn3)-positive β-cells in DKO mice compared to control, suggesting altered Ucn3-Sst signaling; however, only exogenous Sst (Octreotide) and not Ucn3 was able to correct the decreased glucose excursion.

**Conclusions:** Loss or inhibition of both Casp1 and Ripk3 fundamentally alter islet responses to glucose. Our findings highlight that endogenous Casp1 and Ripk3 act independently of inflammatory or cell death signals to coordinate normal glucose-stimulated insulin release.

## Introduction

The global prevalence of overweight and obesity continue to rise, an observation heavily influenced by genetic predisposition, physical inactivity, energy-rich and over-processed diets and socio-economic factors [1]. In people living with obesity, metabolic dysfunction links nutrient-overload and chronic low-grade inflammation to insulin resistance, type 2 diabetes (T2D), metabolic dysfunction-associated fatty liver disease and cardiovascular disease [2; 3]. Importantly, inflammation associated with the progression of metabolic disease can instigate cell death programs across an array of metabolic tissues, including the endocrine pancreas [4–8]. Recent studies have furthered our knowledge of how inflammatory and cell survival pathways are engaged upon nutrient stress [9; 10]. However, there remains a gap in our understanding as to the role of these programs in the maintenance of metabolic homeostasis across the progression of diet-induced obesity.

Inflammatory programs and if required, cell death pathways, are necessary processes in all cells. Moreover, immune and metabolic cues allow for maintenance or the return to cellular homeostasis. Caspase-1 (Casp1) is an enzyme that is positioned at a nexus of inflammation and cell death [11]. Arguably best known for its functions in processing pro-interleukin (IL)-1β and pro-IL-18 via canonical inflammasome activation, Casp1 can also signal to induce an inflammatory form of cell death called pyroptosis via cleavage of Gasdermin D (GSDMD). Caspase 11 (Caspase 4/5 in humans) also cleaves GSDMD via a non-canonical pathway. Genetic disruption of Casp1 in preclinical settings has not provided clarity as to the role of this important regulator in models of diet-induced metabolic dysfunction [12–16]; however, tangible beneficial effects were observed on T2D progression [4; 17–20]. Moreover, Casp11 activity increases during nutrient stress and Casp11 deletion proved protective against obesity-induced metabolic dysfunction [21–23].

Necroptosis is a programmed form of necrotic cell death unlike apoptosis (programmed and non-inflammatory) or necrosis (not programmed and inflammatory). The necroptotic pathway is mediated by the actions of two kinases, receptor-interacting serine/threonine-protein kinase 1 (Ripk1), Ripk3 and an effector protein, mixed lineage kinase domain-like (Mlkl), which acts as the final step in the pathway and ruptures the cell membrane from within [11]. In the context of metabolic disease, the disruption of necroptosis via deletion or therapeutic targeting is widely recognized as beneficial, though genetic studies have focused on mutant versions of Ripk1 due to its essential role in development, or on disruption of Ripk3 [24–28]. While inflammation and cell death programs have been the primary focus of most studies, it is now well established that Casp1 and Ripk3 can directly regulate multiple mitochondrial and metabolic targets [29–32].

Given the interplay between inflammation, cell death, and metabolic homeostasis, we sought to understand the effect of dual disruption of Casp1 and Ripk3 on aspects of whole-body metabolism in mice when fed a high fat diet (HFD) or control, matched low-fat diet (LFD). We hypothesized that blunting Casp1-mediated inflammation and Ripk3-mediated necroptotic signaling would improve metabolic outcomes during diet-induced obesity. Using a series of genetic loss of function models, *in vivo* metabolic assessments, and *ex vivo* assays, we demonstrate that Casp1 and Ripk3 function is required for appropriate glucose-stimulated insulin secretion (GSIS) through modulation of the urocortin-3 (Ucn3)-somatostatin (Sst) feedback loop in pancreatic islets. Together, this work broadens the conceptual framework as to the regulators of glucose tolerance and insulin secretion.

## Materials and methods

### Mouse models

Mice harbouring a whole-body deletion of Casp1 and Casp11, generated by Dr. Richard Flavell [33], were crossed with mice harbouring a whole-body deletion of Ripk3, generated by Dr. Vishva Dixit [34]. Double heterozygous mice were crossed to produce littermate-derived WT, Casp1/11 KO, Ripk3 KO and double Casp1/11/Ripk3 mice (DKO). To interrogate the relative contributions of the Casp1/11 dual knockout, we also crossed B6.Cg-Casp1^em1Vnce^/J [35] (JAX#032662) with Ripk3 KO mice to rederived WT, single and DKO animals with intact Casp11 signaling. For pharmacological studies, we used C57BL/6J mice, originally purchased from The Jackson Laboratory (JAX#00664), bred in house and backcrossed yearly.

All mice were maintained on a C57BL/6J background. Male and female mice were reared on a standard laboratory chow diet (58 % kcal from carbohydrates, 18 % kcal from fat, and 24 % kcal from crude protein; Harlan Teklad #2018) until 8 weeks of age. At this point, male and female mice were placed on either a standardized low-fat diet (LFD) (10% kcal from fat, 70% kcal from carbohydrates and 20% kcal from protein; Research Diets Inc, #D12450B) or a high fat diet (HFD) (∼60% kcal from fat, ∼20% kcal from carbohydrates and ∼20% kcal from protein; Research Diets Inc, #D12492) for 14-weeks. Mice were group-housed in ventilated cages at ∼22°C, maintained on a 12-hour light/dark cycle (lights on at 07:00) with unrestricted access to food and water. Mice were euthanized in the fed state following administration of ketamine (150 mg/kg)-xylazine (10 mg/kg) anesthesia, followed by cervical dislocation. All animal experiments were performed in accordance with the guidelines of the Canadian Council on Animal Care and were approved by the University of Ottawa Animal Care Committee (BMIe3364).

### Metabolic assessments

Body weights were recorded weekly throughout the 14-week dietary intervention. Body composition was assessed using an EchoMRI-700 (EchoMRI) to determine lean and fat mass. Mice were singly housed and acclimated for 24 h before undergoing indirect calorimetry assessment in a Comprehensive Laboratory Animal Monitoring System (CLAMS; Columbus Instruments) for a further 72 h. Following data output from the Oxymax software, comparisons between groups of mice were conducted using CalR 2.0 [36].

Blood glucose levels were measured using a handheld glucometer (Accuchek), following introduction of a small cut in the lateral tail vein. Intraperitoneal (IP) glucose tolerance test (ipGTT), or oral (oGTT) were conducted following a 4 h fast, beginning at ∼9 am. Following basal glucose measurement, mice received 1.5 g/kg (ipGTT) or 2 g/kg (oGTT) D-glucose in saline. Blood glucose concentrations were subsequently measured at 20, 40, 60, 90, and 120 min, with blood samples collected in lithium heparin-coated tubes at baseline and 20 min for plasma insulin measurements. Following blood collection, samples were left to clot at room temperature for approximately 30 min and then centrifuged at 11,000 x g for 5 min. The resulting supernatant was collected, and insulin concentrations were measured by ELISA (ALPCO, #80-INSMSU-E10).

For experiments involving Casp1 and Ripk3 pharmacological inhibition, C57BL/6J mice were administered the Casp1 inhibitor Ac-YVAD-cmk (2.5 mg/kg; Cayman Chemical; #10014) and the Ripk3 inhibitor GSK’872 (2.5 mg/kg; Cayman Chemical; #23300) by IP injection once daily for two consecutive days (8:00 am). Glucose tolerance was assessed by ipGTT on day 2. For Ucn3 experiments, mice were given an IP injection with a Ucn3 peptide (Bachem; 4040194) 5 min prior to glucose administration at the indicated concentrations. For Octreotide injections to mimic somatostatin, Octreotide (25 µg/kg) was injected (IP) 30 min prior to glucose administration.

For insulin tolerance tests (ITT), mice were fasted for 4 h in the morning (9:00 am) and basal blood glucose was assessed prior to IP injection of insulin (0.5 U/kg) in saline (NovoRapid^®^). Blood glucose concentrations were measured at 20, 40-, 60-, 90-, and 120-min following insulin administration.

### Islet isolation

Mouse pancreatic islets were isolated as previously described [37]. Briefly, the pancreatic duct was perfused with a collagenase solution (1,000 U/mL; Sigma-Aldrich, #C7657) prepared in Hank’s Balanced Salt Solution (HBSS; 137 mM NaCl, 5.4 mM KCl, 4.2 mM NaH_2_PO_4_, 4.1 mM KH_2_PO_4_, 10 mM HEPES, 1 mM MgCl_2_, 5 mM dextrose; pH 7.2) following clamping of the duct at the duodenum. Following ductal inflation, the pancreas was excised and transferred to conical tubes containing ice-cold HBSS. Samples were digested in a 37°C water bath for 10.5 min, after which tubes were vigorously shaken 10-15 times to further dissociate pancreatic tissue. Digestion was immediately quenched by the addition of ice-cold HBSS supplemented with CaCl_2_ (1 mM). Samples were washed three times in HBSS + CaCl_2_ with centrifuging in between for 1 min at 1000 *x g* and, after the final wash, tissue was resuspended in RPMI medium (Multicell; #350-000 CL) supplemented with 1% penicillin-streptomycin (Gibco; #15140122) and 10% fetal bovine serum (Multicell; #090-150). Islets were separated from exocrine tissue using a Histopaque density gradient (Sigma-Aldrich; #10771) and collected using a 70 μm cell strainer. Islets were then hand-picked to ensure purity using a dissecting microscope and cultured overnight at 37°C in supplemented RPMI in a 5% CO_2_ incubator.

### Immunofluorescence staining and image quantification

Whole pancreata were harvested from male and female DKO mice and their littermate-derived WT controls following LFD or HFD feeding. Pancreata were fixed in 10% neutral-buffered formalin and processed for paraffin embedding through the Louise Pelletier Histology Core (University of Ottawa). Paraffin-embedded tissue was sectioned at 5 μm thickness using a Leica RM2135 microtome and mounted onto glass slides. Immunofluorescence (IF) staining was conducted as previously described [38]. To summarize, tissue sections were first deparaffinized and rehydrated through a series of graded xylene and ethanol washes, followed by a phosphate-buffered saline (PBS) rinse. Heat-induced epitope retrieval was performed using an EZ Retriever microwave in citrate buffer (10 mM sodium citrate, 0.05% Tween-20; pH 6.0) at 95°C for 10 min. A hydrophobic barrier was created around each tissue section using an ImmEdge Pen (Vector Laboratories; #H-4000), and DAKO serum-free protein block (Agilent; #X090930-2) was applied for 30 min in a humid chamber. After removing the protein block, primary antibodies were added to the sections and incubated overnight at 4°C. The following day, slides were washed three times with PBS, then incubated with Alexa Fluor-conjugated secondary antibodies for 1 hour at room temperature in a humid chamber. After three additional PBS washes, slides were mounted with Vectashield HardSet mounting medium with DAPI (Vector Laboratories; #H-1500) and left to harden under a coverslip. Primary antibodies were all diluted in Dako Antibody Diluent solution (Agilent; #S202230-2). The following primary antibodies were used: rabbit anti-somatostatin (1:500, Sigma-Aldrich; #HPA019472), mouse anti-insulin (1:250, Cell Signaling; #8138S), mouse anti-glucagon (1:250, Sigma-Aldrich; #G2654-0.2ml), rabbit anti-insulin (1:200, Cell Signaling; #3014), rabbit anti-Ucn3 (1:500, custom antibody generated at the Salk Institute). The following secondary antibodies were used: goat anti-rabbit immunoglobulin G (IgG) (H+L), Alexa Fluor 594 (1:1000, Life Technologies; #A11037) and goat anti-mouse IgG (H+L), Alexa Fluor 488 (1:1000, Life Technologies; #A11029).

For quantification of islet cellular composition, pancreatic sections were scanned using a Zeiss Axio Scan.Z1 slide scanner at the Louise Pelletier Histology Core (University of Ottawa). For each mouse, all islets containing ≥30 cells were analyzed by a single investigator who was blinded to sex and genotype during image analysis. Between 5 and 59 islets per sample were manually quantified based on immunofluorescence area using QuPath v0.5.0 bioimaging analysis software. For each biological replicate, the mean of all islet measurements was used for statistical analysis and represented by the larger data points, while individual islet values are also shown in the background of each plot. The percentage of insulin-, somatostatin-, and glucagon-positive area was calculated as [(hormone area / total islet area) × 100]. The percentage of urocortin 3-positive area was calculated as [(urocortin 3 area / insulin area) × 100].

### Dynamic glucose-stimulated insulin secretion assay

Dynamic GSIS was assessed in islets isolated from DKO mice and littermate-derived WT controls, as well as in islets from C57BL/6J mice treated *in vitro* with either DMSO (vehicle control) or a combination of Ac-YVAD-cmk (100 µM) and GSK’872 (5 µM) for 24 h. Islets were assayed using a Biorep Technologies system. For each mouse, 70 islets were handpicked and loaded into Perspex microcolumns between two layers of acrylamide-based microbeads (Biorep Technologies; PERI-BEADS) and exposed to various glucose concentrations diluted in Krebs-Ringer bicarbonate HEPES buffer (KRBH; 115 mM NaCl, 5 mM KCl, 24 mM NaHCO_3_, 2.5 mM CaCl_2_, 1 mM MgCl_2_, 10 mM HEPES, 0.1% w/v BSA). Following a 40 min pre-incubation in KRBH containing low glucose (LG) at a flow rate of 40 µL/min, islets were sequentially exposed to KRBH containing varying glucose concentrations as follows: LG (2.8 mM) for 15 min at 40 µL/min; high glucose (HG; 16.7 mM) for 15 min at 80 µL/min; HG for 30 min at 40 µL/min; LG for 25 min at 40 µL/min; KCl (30 mM) for 15 min at 80 µL/min; KCl (30 mM) for 20 min at 40 µL/min; and finally LG for 25 min at 40 µL/min. Perifusate fractions were collected every 2.5 or 5 min and maintained at 4 °C throughout the experiment. Islets were maintained at 37°C throughout the experiment. Samples were stored at −20°C short term until insulin concentrations were quantified by ELISA (ALPCO; #80-INSMR-CH10).

### RNA isolation, cDNA synthesis and qPCR

Following islet isolation, mouse islets from DKO and littermate-derived WT controls were handpicked, lysed in Buffer RLT containing 1% β-mercaptoethanol and stored at −80°C until processing. Total RNA was extracted using TriPure RNA Isolation Reagent (Roche; #11667165001) according to the manufacturer’s instructions. RNA concentrations were measured using a Take3 Plate (Agilent; #TAKE3-SN) on a Synergy H1 plate reader (BioTek) and normalized with DNase/RNase-free water prior to transfer into 8-strip PCR tubes. Reverse transcription was performed by adding All-in-One 5× RT Master Mix (Applied Biological Materials; #G592) to each sample, followed by cDNA synthesis according to the manufacturer’s protocol. The resulting cDNA was diluted 1:10-1:40, depending on gene target in DNase/RNase-free water. qPCR was conducted using QuantiNova Probe PCR Master Mix (Qiagen; #208254) for TaqMan probes, while BlasTaq 2X qPCR Master Mix (Applied Biological Materials; #G891) was used for non-TaqMan probes. Primer sequences are provided in Supplementary Table X. Reactions were run on a Rotor-Gene Q system (Qiagen). Relative transcript expression was calculated using the 2^-ΔΔCt^ [39] method with *Ppia* and *Hprt* as housekeeping genes.

### Statistical Analysis

All statistical analyses were performed using GraphPad Prism (version 10.6, GraphPad Software Inc., La Jolla, CA, USA). Normality of data was determined using the Shapiro-Wilk test (= 0.05). For normally distributed data, differences between groups with a single variable were determined using an unpaired t-test with Welch’s correction. For comparisons involving three or more groups, a one-way ANOVA was used, while a two-way ANOVA was applied when analyzing data with two independent variables. For non-normally distributed data, differences between groups were assessed using an unpaired Mann-Whitney U test. Specific statistical tests are indicated in the corresponding figure legends. Statistical significance was defined as \**p* < 0.05, \*\**p* < 0.01, \*\*\**p* < 0.001, and \*\*\*\**p* < 0.0001. Data are presented as mean ± SEM.

## Results

### Single or dual deletion of Casp1/11 and Ripk3 exacerbates HFD-induced obesity and glucose intolerance in male but not female mice

To test our initial hypothesis that simultaneous disruption of Casp1 and Ripk3 signaling would improve metabolic outcomes during diet-induced obesity, we used mice deficient for Casp1 to disrupt apoptotic inflammatory signaling and mice deficient for Ripk3 to probe the importance of necroptotic pathways. Upon generation of the established Casp1 KO model, the Casp11 gene was also disrupted in *cis*, thereby generating a model deficient for Casp1 and Casp11 [33]. We then crossed Casp1/11 KO mice with Ripk3 KO mice to produce littermate-derived WT, Casp1/11 KO, Ripk3 KO and Casp1/11/Ripk3 DKO lines. Male and female mice of all genotypes were maintained on a chow diet for 8 weeks, followed by a 14-week dietary intervention to a HFD (60% kcal from fat). Subsequently, the mice underwent an array of metabolic phenotyping assessments. (Fig. 1A).

**Fig. 1.**
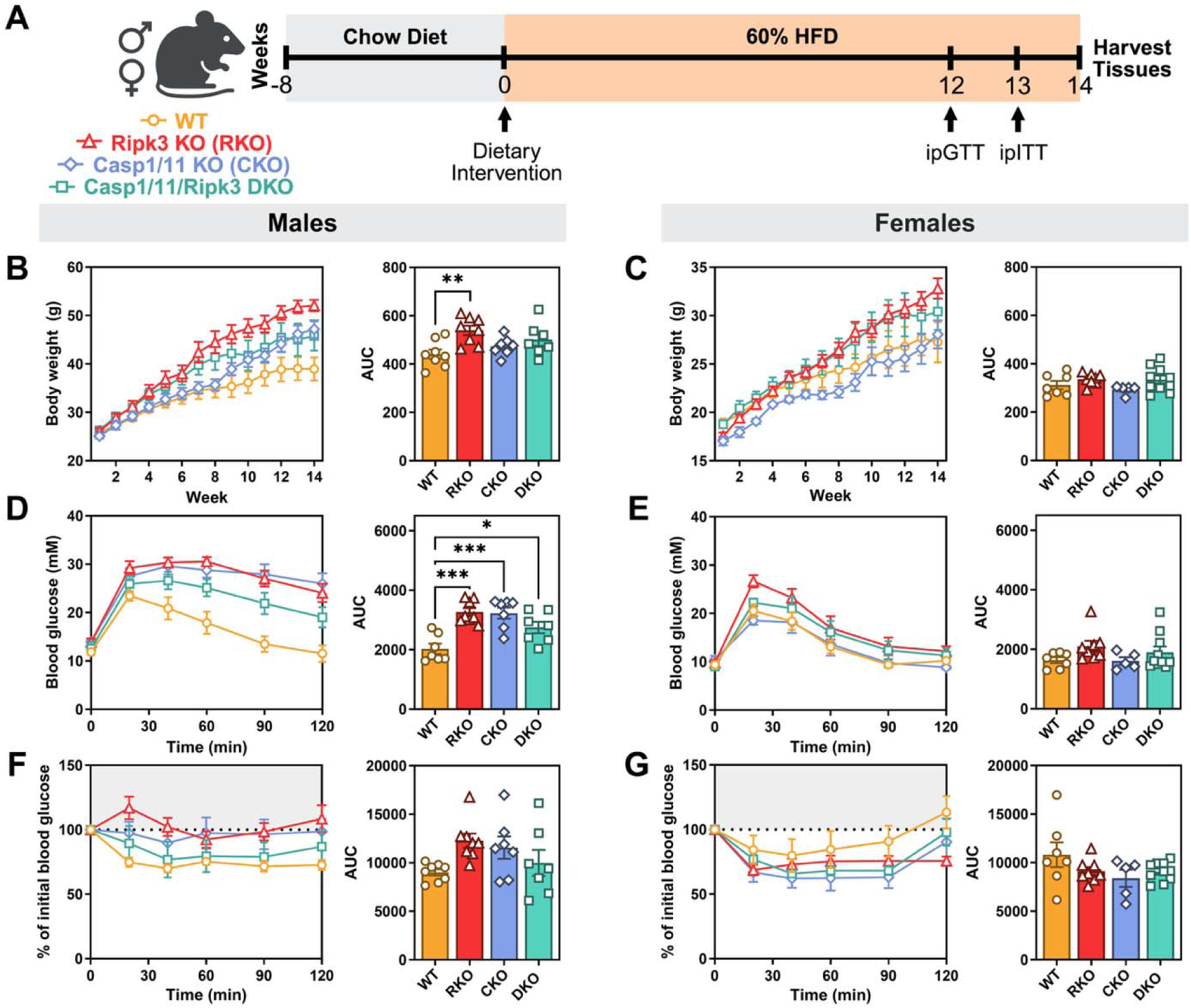
Single or dual deletion of Casp1/11 and Ripk3 exacerbates HFD-induced obesity and glucose intolerance in male but not female mice. (A) Experimental timeline for male and female mice fed a HFD (60% kcal from fat) for 14 weeks. (B, C) Body weights of WT, Ripk3 KO (RKO), Casp1/11 KO (CKO), and Casp1/11/Ripk3 (DKO) mice were measured over 14 weeks. (D, E) ipGTT (1.5 g/kg glucose in saline) was performed on 5-hour fasted mice. Blood glucose levels were measured over 120 min. (F, G) ipITT (1 U/kg insulin) on 4.5-hour fasted mice, with glucose measurements taken over 120 min. Data represents SEM (n = 5-10 mice per genotype). Statistical significance was determined by one-way ANOVA with Tukey’s multiple comparison test. \**P* < 0.05, \*\**P* < 0.01, \*\*\**P* < 0.001.

Although there were no differences in body weight between genotypes at baseline, over the course of several weeks on the HFD, Casp1/11 KO, Ripk3 KO and Casp1/11/Ripk3 DKO male mice began to gain more weight than their WT controls, with body weight differences emerging over the course of the 14-week intervention (Fig. 1B). In contrast, there were no significant differences in female weight gain between genotypes (Fig. 1C). There were no significant differences in the amount of lean or fat mass between genotypes, nor in markers of whole-body energy metabolism measured by metabolic cages between any of the genotypes in either sex (Fig. S1).

To monitor glucose tolerance independent of the gut-assisted incretin response, mice were subjected to an ipGTT 12 weeks after starting the HFD. Congruent with the overall weight gain in male Casp1/11/Ripk3 DKO and single KO mice, glucose tolerance was impaired compared to WT control (Fig. 1D). In comparison, female Casp1/11/Ripk3 DKO and single Ripk3 KO mice showed a trending glucose intolerance (Fig. 1E). Assessment of insulin sensitivity by ipITT revealed a trend towards higher blood glucose levels following insulin administration in male Casp1/11 and Ripk3-deficiencent mice, suggesting reduced insulin sensitivity but no significant differences between genotypes in either sex overall (Fig. 1F and G).

We assessed inflammatory transcript expression in the white adipose tissue (WAT) of male and female control and Casp1/11/Ripk3 DKO mice with no differences (Fig. S2). In the liver of HFD-fed male mice, *Il6* transcript expression was lower and lipid synthesis (*Fasn* and *Srebf1c* transcripts were higher in Casp1/11/Ripk3 DKO compared to WT control (Fig. S2), while no differences were observed in females. Circulating cytokines were also not different between Casp1/11/Ripk3 DKO and WT mice, except for IL-4 in females (Fig. S3). Collectively, these findings indicate a sex-specific propensity for weight gain in male mice when Casp1/11, Ripk3, or both are disrupted. While male mice lacking either Casp1/11 or Ripk3 exhibited impaired glucose tolerance and mildly decreased insulin sensitivity, dual disruption of both genes, did not result in additive or synergistic effects on diet-induced metabolic outcomes.

### Dual deletion of Casp1/11 and Ripk3 improves glucose tolerance without impacting other metabolic measures in lean mice

We next sought to generate a baseline understanding of the metabolic phenotype associated with Casp1/11/Ripk3 genetic deletions in LFD-fed mice. To this end, 8-week-old male and female mice from all genotypes were fed a LFD for 14-weeks and subjected to metabolic phenotyping (Fig. 2A). No differences in body weight (Fig. 2B and C), or lean and fat mass proportions (Fig. S4), or markers of whole-body energy metabolism as assessed by indirect calorimetry in metabolic cages (Fig. S4) were observed between genotypes in either sex. Both the single Casp1/11 and Ripk3 KO mice exhibited glucose tolerance comparable to WT controls. In contrast, both male and female Casp1/11/Ripk3 DKO mice displayed significantly lower glucose excursion curves, suggestive of improved glucose tolerance (Fig. 2D and E). Insulin sensitivity was not different in any genotype of either sex (Fig. 2F and G).

**Fig. 2.**
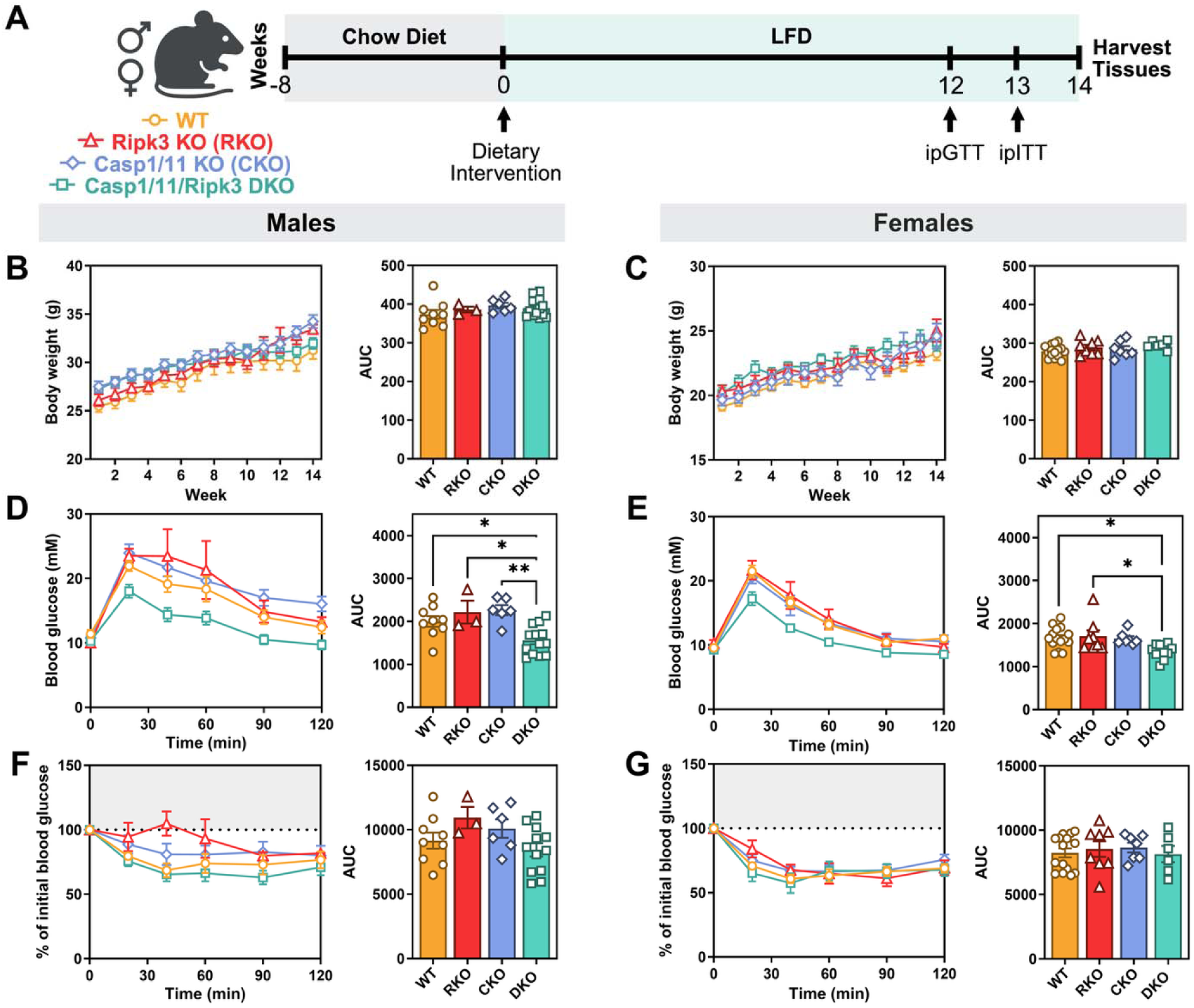
Dual deletion of Casp1/11 and Ripk3 improves glucose tolerance without impacting other metabolic measures in lean mice. (A) Experimental timeline for male and female mice fed a LFD (10% kcal from fat) for 14 weeks. (B, C) Body weights of WT, Ripk3 KO (RKO), Casp1/11 KO (CKO), and Casp1/11/Ripk3 (DKO) mice were measured over 14 weeks. (D, E) ipGTT (1.5 g/kg glucose in saline) was performed on 5-hour fasted mice. Blood glucose levels were measured over 120 min. (F, G) ipITT (0.5 U/kg insulin) on 4.5-hour fasted mice, with glucose measurements taken over 120 min. Data represents SEM (n = 3-15 mice per genotype). Statistical significance was determined by one-way ANOVA with Tukey’s multiple comparison test. \**P* < 0.05, \*\**P* < 0.01.

There were no changes in lipid-related transcript expression in the livers of Casp1/11/Ripk3 DKO mice, and no changes in levels of inflammatory transcripts in the livers or WAT of Casp1/11/Ripk3 DKO mice fed a LFD (Fig. S5). Moreover, circulating cytokines were largely comparable across genotypes in both sexes, with the exception of reduced IL-1β levels in female Casp1/11/Ripk3 DKO mice (Fig. S6). Interestingly, these findings demonstrate that independent of changes in adiposity, weight or insulin sensitivity, and only when both Casp1/11 and Ripk3 are deleted, male and female mice experience enhanced capacity to clear circulating glucose.

### Casp1/11/Ripk3 DKO mice display increased glucose-stimulate insulin secretion in response to glucose

Given the enhanced glucose clearance observed in lean male and female Casp1/11/Ripk3 DKO mice, we examined serum glucose after ∼9 h of overnight fasting (21 h-0800 h) and random-fed insulin levels, measured at ∼0800 h when given unrestricted access to food. Fasting serum glucose was significantly reduced in both male and female Casp1/11/Ripk3 DKO mice (Fig. S7A and C), while insulin levels were significantly higher in female Casp1/11/Ripk3 DKO compared to WT, but not in males (Fig. S7B and D). To assess the *in vivo* plasma insulin levels in response to glucose, blood samples were collected from fasted mice at baseline and 30 min following glucose administration (ip). Plasma insulin levels were significantly elevated in Cap1/11/Ripk3 DKO mice compared to WT controls in both sexes (Fig. 3A). To determine whether this phenotype was driven by intrinsic changes in islet function, we performed dynamic insulin secretion assays *ex vivo* using isolated pancreatic islets. In line with the *in vivo* data, islets from Casp1/11/Ripk3 DKO male and female mice had significantly increased glucose and KCl-simulated insulin secretion (Fig. 3B), demonstrating an enhanced and islet-autonomous effect of dual Casp1/11 and Ripk3 deficiency on insulin secretion. To determine whether this phenotype was maintained under HFD conditions, we next examined insulin secretion in islets isolated from HFD-fed mice. No differences in GSIS or KCl-stimulated insulin secretion were observed in male islets, though a slight but significant increase in insulin secretion remained in islets isolated from females (Fig. S8). These findings suggest that HFD feeding blunts the enhanced insulin secretory phenotype observed under LFD conditions in males and that HFD-induced insulin resistance potentially explains the lack of change in glucose responsiveness in female HFD-fed Casp1/11/Ripk3 DKO mice.

**Fig. 3.**
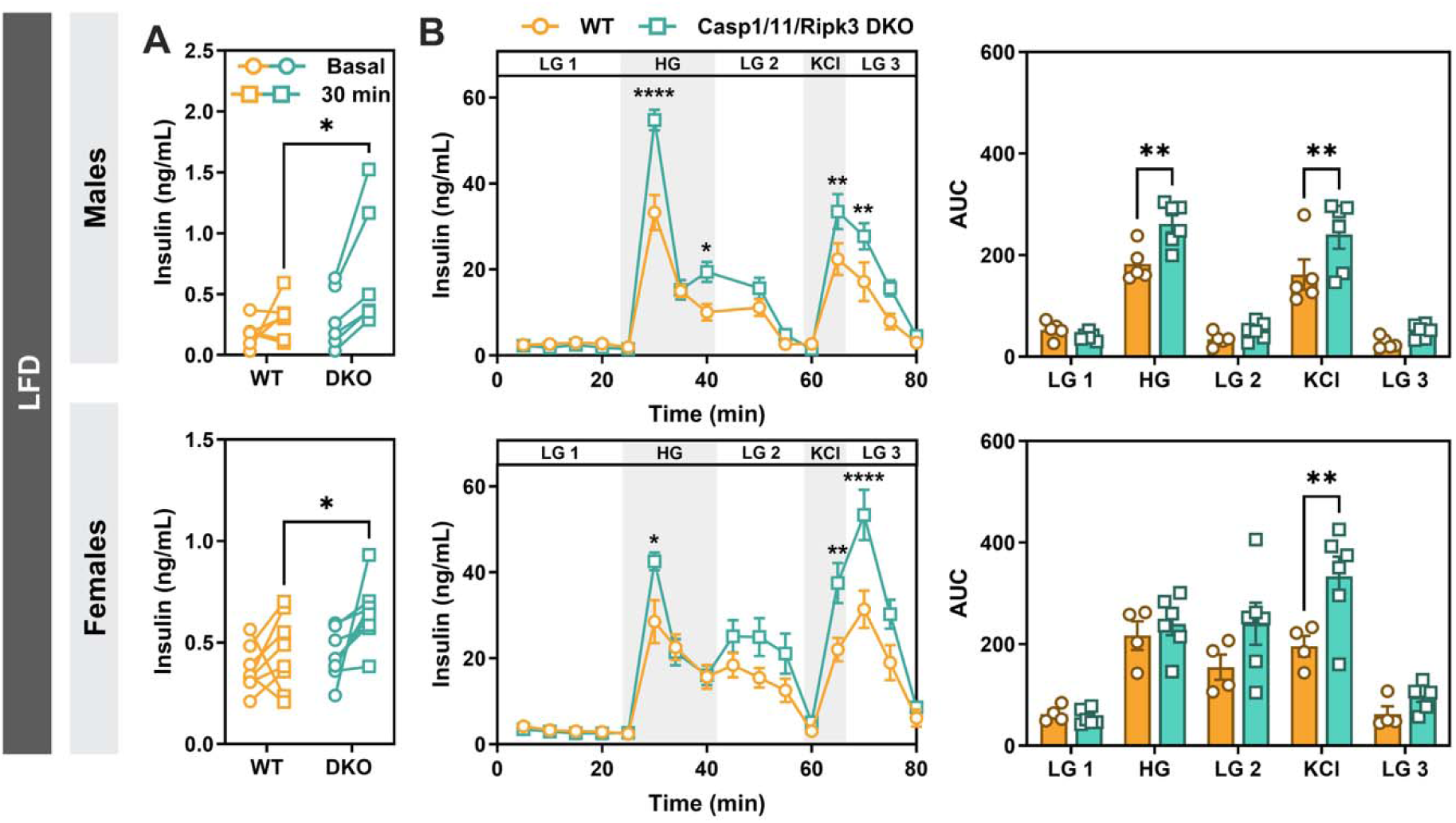
Increased glucose-stimulated insulin secretion in response to glucose in DKO mice. (A) GSIS assessed by measuring serum insulin at baseline and 30 min following IP glucose administration, in male and female WT and DKO mice following ∼12 weeks of LFD feeding. (B) Dynamic insulin secretion was assessed by perifusion of isolated islets from LFD-fed male and female mice, stimulated with low glucose (LG; 2.8 mM), high glucose (HG; 16.7 mM), and depolarizing potassium chloride (KCl; 30 mM). Insulin release was measured over 80 min. Data represents SEM (n = 4-15 mice per genotype). Statistical significance was determined by: (A) two-way repeated-measures ANOVA followed by Fisher’s LSD multiple comparisons test; (B) two-way repeated-measures ANOVA followed by Šidák’s multiple comparisons test. \**P* < 0.05, \*\**P* < 0.01, \*\*\**P* < 0.001, \*\*\*\**P* < 0.0001.

### Cellular composition of Casp1/11/Ripk3 DKO islets is not altered

Since β-cells are responsible for secreting insulin within pancreatic islets, we next examined the transcript levels of key β-cell genes, including *Ins1, Ins2, Glut2, Ucn3, Gck, MafA,* and *Nkx6.1*, in isolated islets from LFD-fed and HFD-fed WT and Casp1/11/Ripk3 DKO mice. While most genes in DKO male and female LFD mice showed no significant changes, the *Ins1* and *Gck* transcripts were notably increased in Casp1/11/Ripk3 DKO males (Fig. 4A and B). In contrast, the HFD Casp1/11/Ripk3 DKO mice had significant transcript increases in *Ins1, Ins2, Glut2, Ucn3* (males), and *Nkx6.1* (males) (Fig. S9). To determine whether other endocrine cell populations were affected, transcripts beyond β-cell identity were also assessed. In the LFD group, no differences were observed in *Crhr2*, the δ-cell receptor responsible for binding Ucn3 (Fig. 4A and B). Additionally, transcripts for glucagon (*Gcg*) and somatostatin (*Sst*) were measured in the HFD group, with no significant differences detected between genotypes (Fig. S9).

**Fig. 4.**
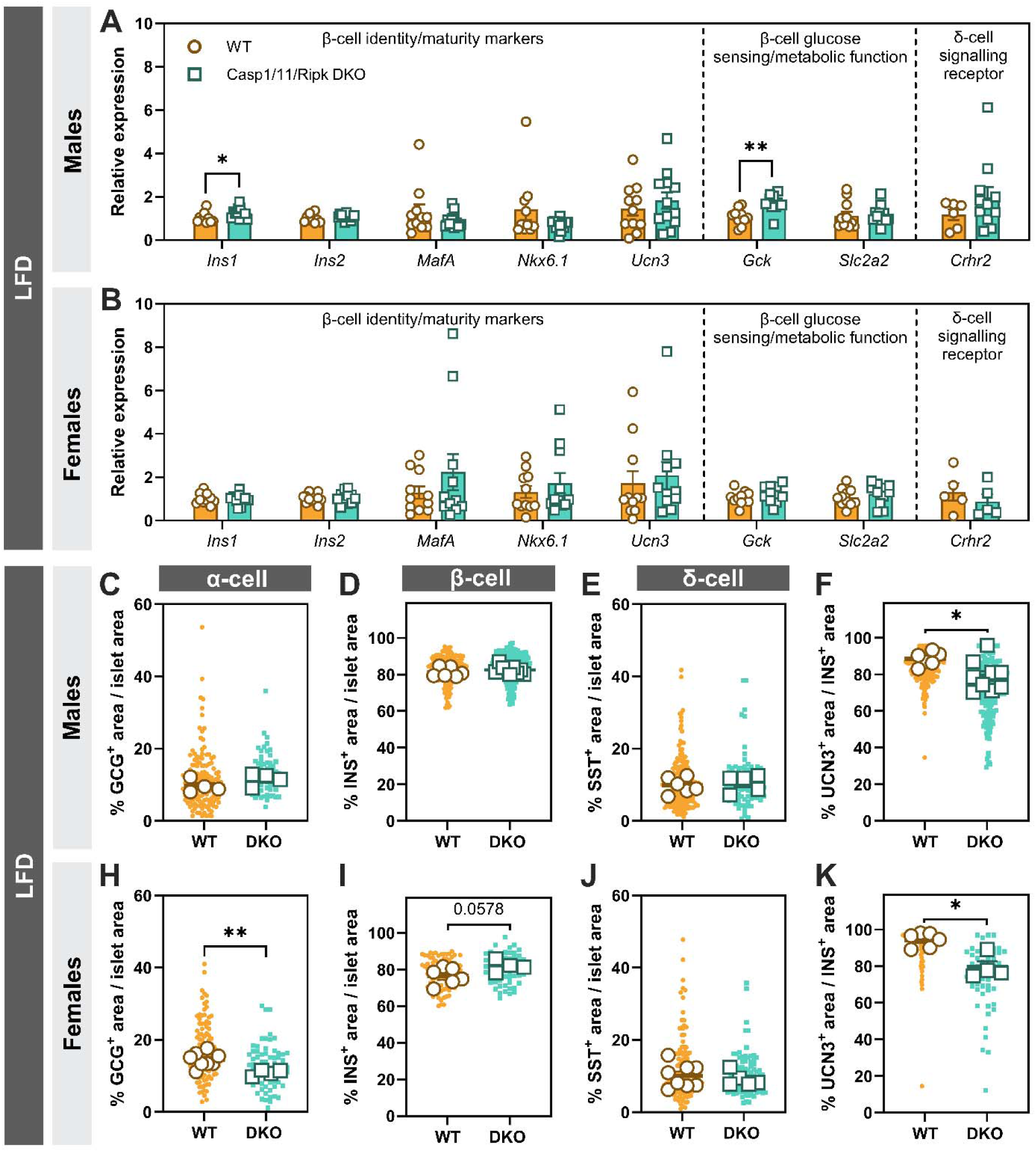
Cellular composition of Casp1/11/Ripk3 DKO islets was not significantly altered. (A, B) Relative transcript expression of genes associated with β-cell identity and maturity (*Ins1, Ins2, MafA, Nkx6.1, Ucn3*), β-cell glucose sensing and metabolic function (*Gck, Slc2a2*), and δ-cell signalling (*Crhr2*) was measured in isolated islets from male and female mice maintained on a LFD. Relative transcript abundance was quantified by RT-qPCR. Data are presented as mean ± SEM (n = 5-12 mice per genotype). (C-K) Paraffin-embedded pancreatic sections from male and female mice fed a LFD were analyzed by IF staining to quantify selected islet hormones. (C, H) % glucagon (Gcg) area normalized to the total area of each islet to represent the percent α-cells. (D, I) % insulin (Ins) area normalized to the total area of each islet to represent the percent β-cells. Representative quantification plots from Ins/urocortin 3 (Ucn3) IF staining are shown. (E, J) % somatostatin (Sst) area normalized to the total area of each islet to represent the percent δ-cells. (F, K) % Ucn3 area normalized to Ins area per islet. Quantification of Gcg, Ins, and Sst area was calculated as (hormone area / total islet area) × 100. The Ins^+^/Ucn3 area was quantified as (UCN3 area / INS area) × 100. Small dots represent individual islets (3-54 per mouse); large dots indicate the mean per mouse (n = 3-9 mice per genotype). Data represents mean ± SEM. Statistical significance was determined by unpaired two-tailed Welch’s t-test (A-K). \**P* < 0.05, \*\**P* < 0.01.

Next, we assess islets cell composition in paraffin embedded pancreatic tissue from male and female mice on either LFD or HFD. The insulin^+^, glucagon^+^ and Sst^+^ area was normalized to the total islet area to quantify the percent β-, α- and δ-cells, respectively. In LFD-fed Casp1/11/Ripk3 DKO male mice, there were no differences in endocrine cell proportions (Fig. 4C-E), while Casp1/11/Ripk3 DKO female mice showed a significant decrease in percent α-cell area, a trending increase in percent β-cell area, and no changes in δ-cell percent area (Fig. 4H-J). Interestingly, under HFD conditions, there was a significant and trending increase in percent β cell area in male and female Casp1/11/Ripk3 DKO islets compared to WT, respectively. This was accompanied by a trending decrease in percent α-cell area and a significant decrease in percent δ-cell area in Casp1/11/Ripk3 DKO islets of both sexes (Fig. S10). To understand how this genotype effect may alter β-cell biology, we assessed the number of β-cells that were Ucn3^+^ and observed a consistent and significant decrease in the number of Ucn3^+^ β-cells in Casp1/11/Ripk3 DKO islets (Fig. 4F and K, Fig. S10). Overall, while the disruption of Casp1/11 and Ripk3 had minimal effect on islet transcript expression or islet endocrine cell composition in LFD-fed mice, the significant decrease in Ucn3^+^ β-cells suggested there could be dysregulated or disrupted intra-islet paracrine action, leading to altered insulin release.

### Increases in insulin secretion in DKO mice are independent of Caspase 11 on a LFD

Casp11 serves a complementary yet distinct inflammatory function to Casp1, including regulation of pyroptotic cell death, cytokine release and organelle trafficking [40]. Given the Casp1/11/Ripk3 mouse model disrupts both Casp1 and Casp11, we next sought to identify whether the deletion of Casp11 is required for the changes in glucose homeostasis we observed. We obtained mice with a 10 bp deletion in the Casp1 gene, but with intact Casp11 function, and crossed these to mice homozygous for Ripk3 deletion to rederive WT, single Casp1 KO, Ripk3 KO and Casp1/Ripk3 DKO.

Following LFD feeding for 14 weeks as previously described above, there were no differences in body weight across genotype in either male or female mice (Fig. S11A and B). Interestingly, glucose excursion during an ipGTT was only significantly lower in Casp1/Ripk3 DKO male mice and displayed a trending decrease in Casp1/Ripk3 DKO female mice (Fig. 5A and B), providing evidence that deletion of Casp1 and not Casp11 was driving the change in glucose homeostasis when Ripk3 is also disrupted. Moreover, when delivered orally, glucose excursion was consistently lower in male Casp1/Ripk3 DKO mice compared to WT (male), providing evidence that genotype differences persist in the presence of incretin-mediated pathways (Fig. S11C). Consistent with previous results, there were no differences in insulin tolerance as assessed by ipITT (Fig. 5C and D). Finally, in this new model of Casp1/Ripk3 deletion, isolated islets secreted more insulin in response to both glucose and KCl during dynamic assessment by perifusion (Fig. 5E), similarly to what was observed in the Casp1/11/Ripk3 DKO model (Fig. 3B). The data highlight that changes to insulin release and glucose homeostasis were mediated by Casp1 even in the presence of Casp11.

**Fig. 5.**
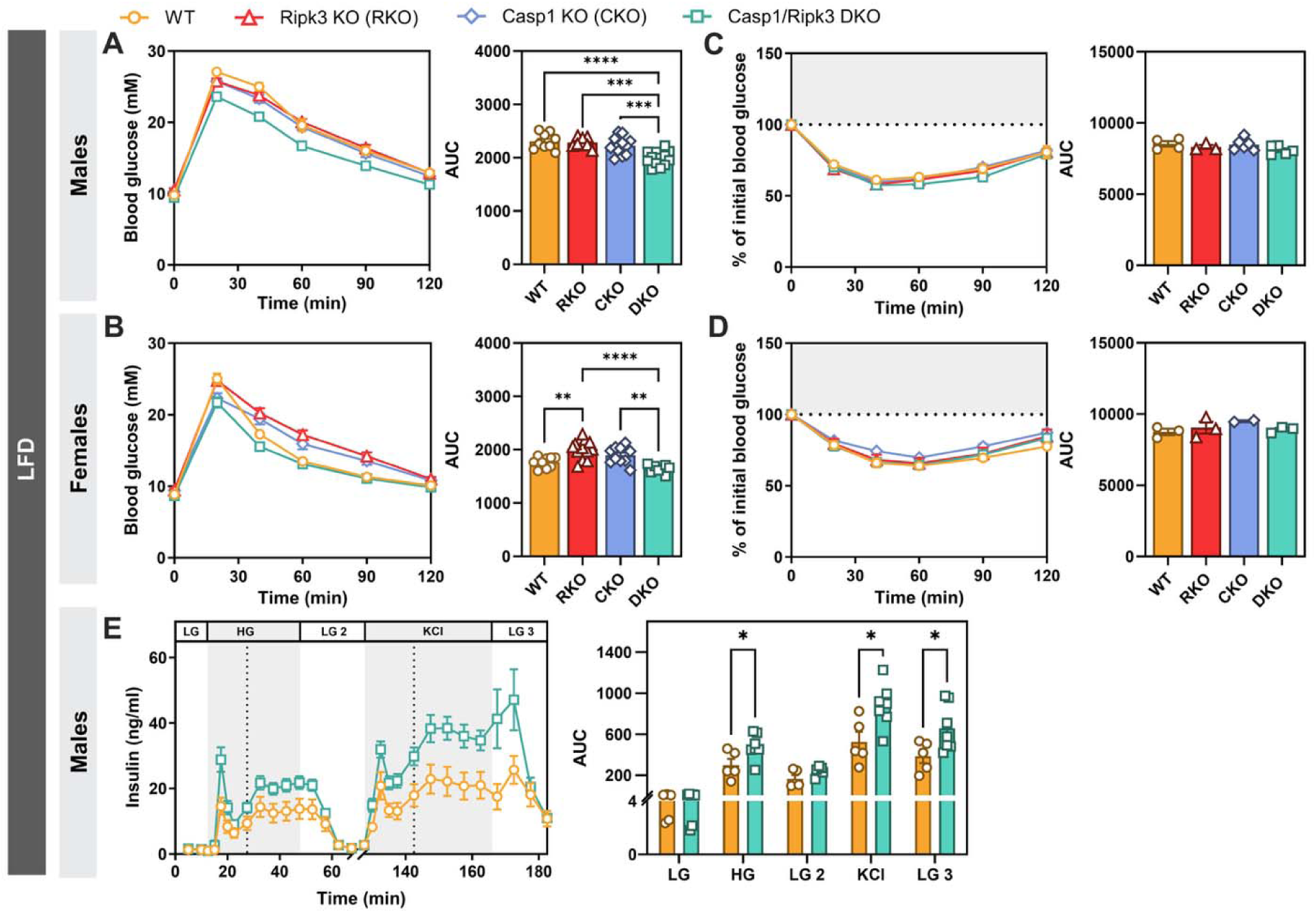
Increases in insulin secretion are independent of Caspase 11. (A, D) Body weight of WT, Ripk3 KO (RKO), Casp1 KO (CKO), and Casp1/Ripk3 (DKO) male and female mice fed a LFD for 14 weeks. (B, E) ipGTT (1.5 g/kg glucose in saline) was performed on 4.5-hour fasted LFD-fed mice. Blood glucose was measured over 120 min. (C, F) ipITT (0.5 U/kg insulin) was performed on 4.5-hour fasted LFD-fed mice, with glucose measured over 120 min. (E) Dynamic insulin secretion was assessed in isolated islets from LFD-fed male WT and DKO mice during perifusion with low glucose (LG; 2.8 mM), high glucose (HG; 16.7 mM), high glucose containing the somatostatin analogue octreotide (HG + Sst; 16.7 mM glucose and 0.5 µM octreotide), and depolarizing KCl (30 mM). Insulin secretion was measured over a 182.5-min perifusion period. Perifusion flow rates were as follows: LG 1-4, 40 µL/min; HG 1, 80 µL/min; HG 2, 40 µL/min; HG + Sst 1, 80 µL/min; HG + Sst 2, 40 µL/min; KCl 1, 80 µL/min; and KCl 2, 40 µL/min. Samples between 65-120 min were quantified but removed due to lack of relevance for the experiment. Data represents mean ± SEM (n = 2-13 mice per genotype). Body weight and GTT/ITT AUC values were analyzed using a one-way ANOVA with Tukey’s multiple comparisons test. \*\**P* < 0.01, \*\*\**P* < 0.001, \*\*\*\**P* < 0.0001.

### Dual pharmacological inhibition of Casp1 and Ripk3 increases glucose-stimulated insulin release and glucose homeostasis in islets and mice

Dual deletion of Casp1 and Ripk3 affected insulin secretion and glucose tolerance in two separate genetic mouse models. To complement this, we next examined whether acute pharmacological inhibition would mirror these phenotypes. We used C57BL/6J male and female mice and administered saline as a vehicle for two consecutive days followed by an ipGTT on day 2 to establish a baseline glucose tolerance. Mice were next administered a combination of the Casp1 inhibitor Ac-YVAD-cmk (2.5 mg/kg) and the Ripk3 inhibitor GSK’872 (2.5 mg/kg) for two consecutive days, with another ipGTT on day four (Fig. 6A).

**Fig. 6.**
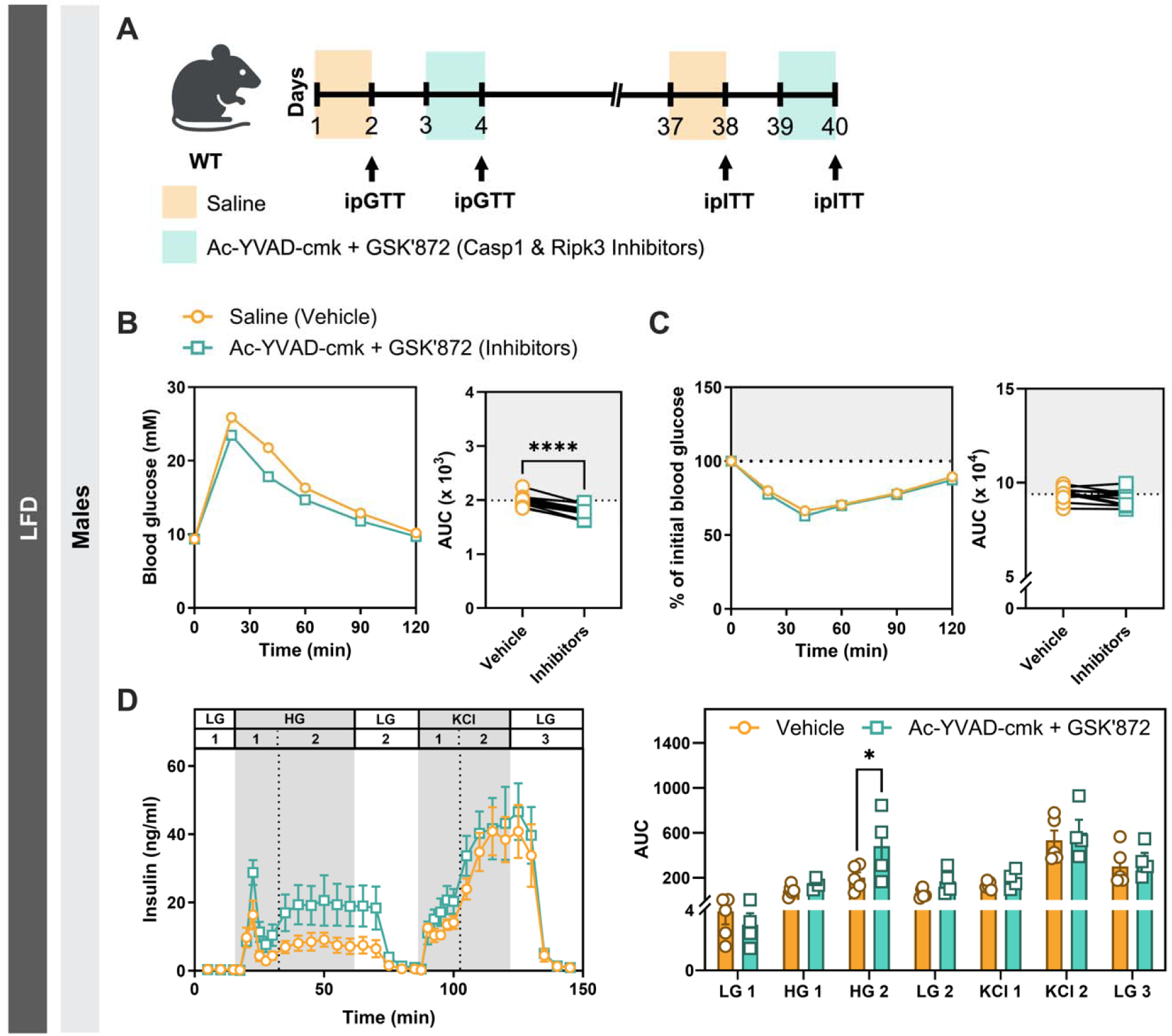
Dual pharmacological inhibition of Casp1 and Ripk3 increases glucose-stimulated insulin release and glucose homeostasis in islets and mice. (A) Experimental timeline for male C57BL/6J mice. Mice received IP injections of saline or Casp1/Ripk3 inhibitors (Ac-YVAD-cmk and GSK’872; 2.5 mg/kg in saline) over 2 consecutive days where each mouse received all treatments longitudinally. (B) IP GTTs (2 g/kg glucose in saline) were performed following a 4.5 h fast, with blood glucose measured over 120 min. (C) IP ITTs (0.6 U/kg insulin) were performed in 4.5 h-fasted mice, with blood glucose measured over 120 min. (D) Dynamic insulin secretion from isolated islets from C57BL/6J male mice treated with either DMSO (vehicle control) or a combination of the Casp1/Ripk3 inhibitors Ac-YVAD-cmk (100 µM) and GSK’872 (5 µM) for 24 h in response to low glucose (LG; 2.8 mM), high glucose (HG; 16.7 mM), and depolarizing potassium chloride (KCl; 30 mM). Insulin secretion was measured over a 150-min perifusion period. Perifusion flow rates were as follows: LG 1-3, 40 µL/min; HG 1, 80 µL/min; HG 2, 40 µL/min; KCl 1, 80 µL/min; and KCl 2, 40 µL/min. Data represents mean ± SEM (n = 4-11 mice per condition). Statistical significance was determined using: (B, C) repeated-measures one-way ANOVA followed by Tukey’s multiple comparisons test, and (D) repeated-measures two-way ANOVA followed by Šidák’s multiple comparisons test. \*\**P* < 0.01, \*\*\*\**P* < 0.0001.

Pharmacological inhibition of Casp1 and Ripk3 significantly lowered glucose excursion, recapitulating the metabolic phenotype observed in the genetic DKO models in both males (Fig. 6B) and females (Fig. S12A). A single dose of both drugs prior to a 4 h fast caused a trending decrease in blood glucose levels 20 min post glucose administration during an ipGTT (Fig. S12B); however, this was not significant and revealed that at least 2 consecutive days of inhibitor delivery was required. Moreover, this effect was lost following a 1-week washout period and was not observed when mice were injected with each compound in isolation (Fig. S12C). There was no change in insulin sensitivity when comparing 2 days of saline and 2 days of Casp1 and Ripk3 inhibitors (Fig. 6C), supporting a pharmacological effect that involves insulin secretion.

To further assess whether acute pharmacological inhibition could reproduce the genetic phenotype *ex vivo*, isolated islets from male C57BL/6J mice were treated with vehicle control or a combination of Ac-YVAD-cmk (100 µM) and GSK’872 (5 µM) for 24 h, followed by dynamic assessment of GSIS by perifusion. Pharmacological inhibition of Casp1 and Ripk3 enhanced GSIS, while KCl-induced insulin release remained unchanged (Fig. 6D). The enhanced GSIS observed following acute pharmacological inhibition suggests that the phenotype in genetic DKO mice is unlikely to result from developmental compensation associated with lifelong genetic deletion.

### Lack of Casp1 and Ripk3 results in altered intra-islet crosstalk

Ucn3 is co-secreted with insulin upon glucose stimulation and plays an important role in stimulating Sst secretion to mediate negative regulation of β-cell insulin release [41]. Given that we identified a significant decrease in the number of Ucn3^+^ β-cells in the LFD-fed male and female Casp1/11/Ripk3 DKO islets, we reasoned that lower Ucn3 secretion may, in turn, lead to a deficient Sst response, which would allow for the increase in GSIS. To assess this, WT and Casp1/Ripk3 DKO mice underwent two ipGTTs separated by 2 days. During the first ipGTT, mice received an IP injection of saline as a vehicle 5 min before glucose administration, while during the second ipGTT mice received recombinant mouse Ucn3 5 min before glucose administration. Thus, the effects of Ucn3 were assessed within the same mouse using the saline-treated ipGTT as the control (Fig. 7A).

**Fig. 7.**
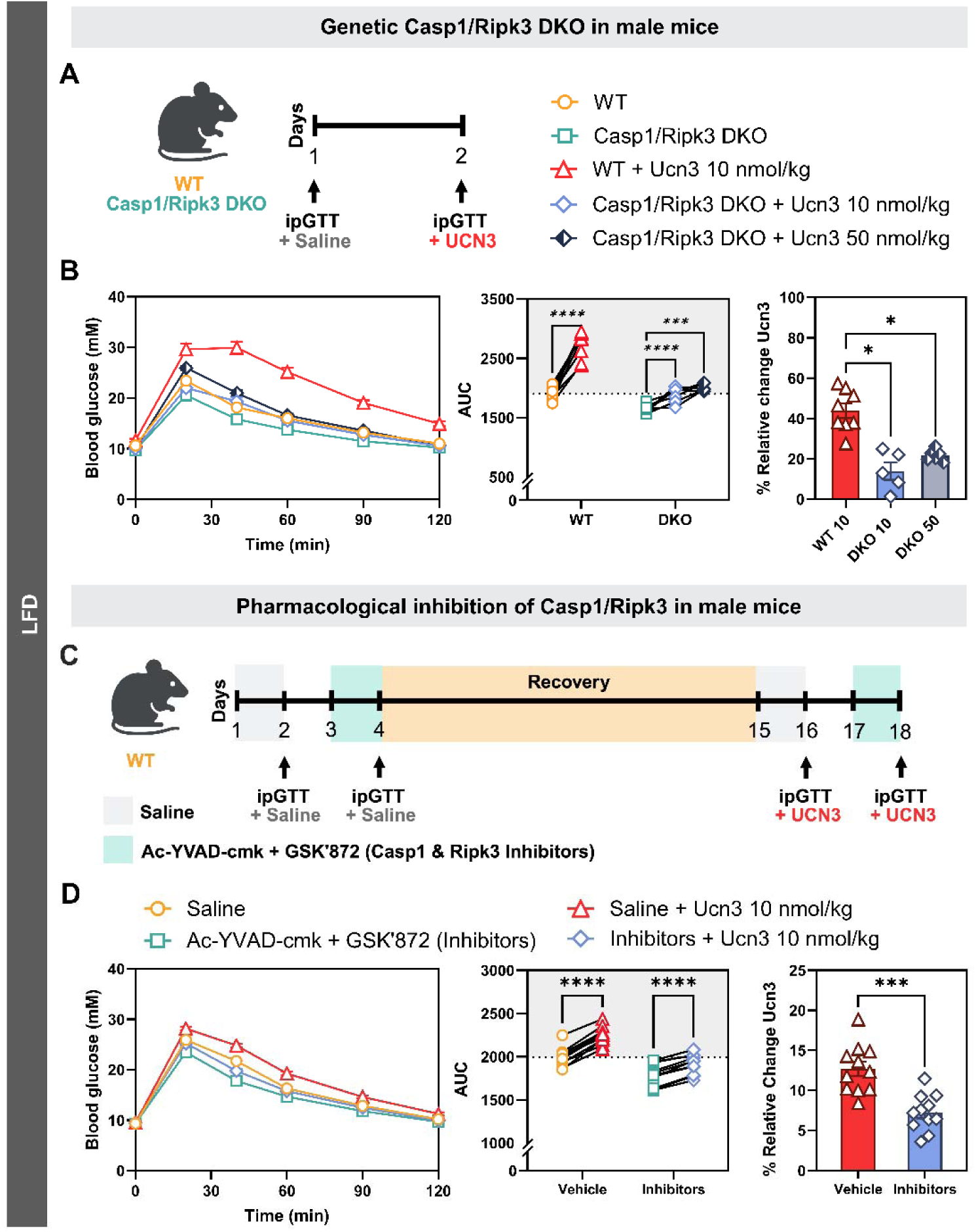
Alterations in glucose homeostasis were not rescued by exogenous Ucn3. (A) Experimental timeline for a longitudinal study of WT and Casp1/Ripk3 DKO mice fed a LFD, with each mouse serving as its own control. (B) Following a 4.5-hour fast, male WT and DKO mice were administered ip injections of saline, followed 2 days later with exogenous Ucn3 (10 or 50 nmol/kg) 5 min before an ipGTT (2 g/kg glucose in saline). Blood glucose was monitored over 120 min. (C) Experimental timeline of C57BL/6J mice for 18 days, with each mouse serving as its own control. (D) C57BL/6J mice received two IP injections of saline or Casp1/Ripk3 inhibitors (Ac-YVAD-cmk and GSK’872; 2.5 mg/kg in saline) over two days. On day 16 and 18, following a 4.5-hour fast, Ucn3 (10 nmol/kg) was administered 5 min prior to an ipGTT (2 g/kg glucose in saline), with blood glucose monitored over 120 min. Paired AUC values were calculated for all conditions, and the percent relative change in response to Ucn3 was calculated as [(Ucn3 AUC - saline AUC) / saline AUC × 100].

In WT mice, Ucn3 led to a robust upward shift in the glucose excursion curve, while the response in the Casp1/Ripk3 DKO mice was significantly blunted. Moreover, even with a dose of Ucn3 5x higher, the Casp1/Ripk3 DKO mice were still able to clear glucose similarly to WT controls (Fig. 7B). In addition, we used C57BL/6J mice that had been injected with either vehicle and then Casp1 and Ripk3 inhibitors with Ucn3 administration prior to the ipGTT, obtaining the same result as the genetic model, where the Ucn3-dependent effect on glucose excursion was significantly higher in vehicle-treated compared to inhibitor-treated mice (Fig. 7C, D and Fig. S13). This strongly suggests that while there may be alterations in the Ucn3 axis, the main mechanistic explanation for increased insulin secretion lies downstream of Ucn3 release.

To assess whether there were defects in Sst signaling, we took a similar approach as above and administered Octreotide, a modified Sst peptide, or saline, 30 min prior to an ipGTT in WT versus Casp1/Ripk3 DKO mice (each mouse received a saline ipGTT and an Octreotide ipGTT, separated by 2 days) (Fig. 8A). Interestingly, while Casp1/Ripk3 DKO mice were resistant to Ucn3-induced glucose intolerance, Octreotide pre-treatment increased the glucose excursion curve in both WT and Casp1/Ripk3 DKO mice to the same extent, in both male and female mice (Fig. 8B and C). Surprisingly, Casp1/Ripk3 DKO islets secreted the same amount of Sst compared to WT islets in response to high glucose, despite higher insulin levels during a perifusion experiment (Fig. S14). These experiments revealed that Casp1/Ripk3 DKO islets retained the ability to transmit the signal downstream of the Sst receptor; however, they suggest that there are defects in β-cell to δ-cell communication when both Casp1 and Ripk3 are deficient or inhibited.

**Fig. 8.**
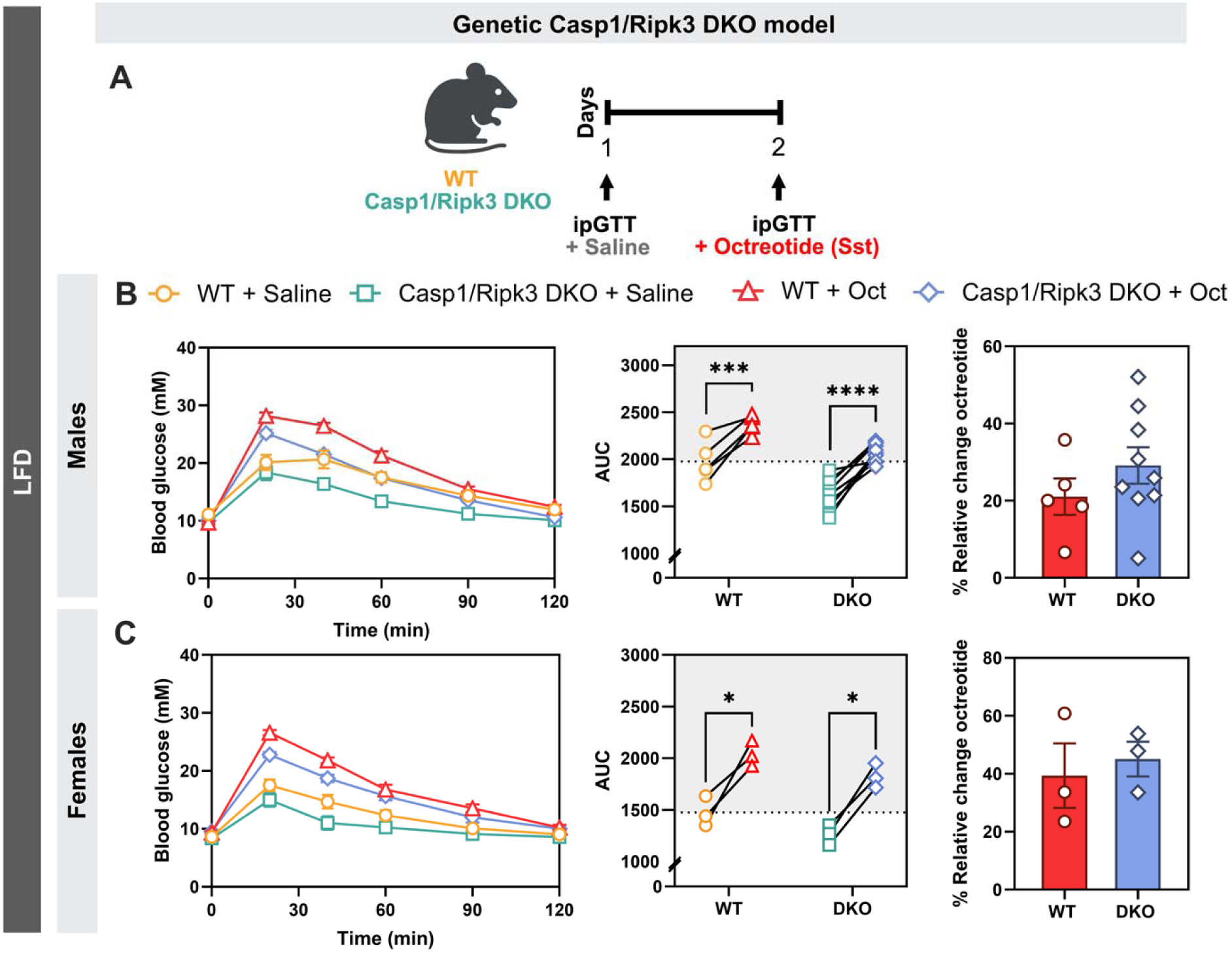
Exogenous Sst corrects the exacerbated glucose response in DKO mice. (A) Experimental timeline of WT and Casp1/Ripk3 DKO mice fed a LFD, with each mouse serving as its own control. (B-C) Following a 4.5-hour fast, male and female WT and DKO mice were administered IP injections of saline, followed 2 days later by the somatostatin analogue octreotide (25 µg/kg) 30 min prior to an ipGTT (2 g/kg glucose in saline). Paired AUC values were calculated, and the relative change in glucose excursion following octreotide treatment was expressed as [(octreotide AUC − saline AUC) / saline AUC × 100]. Data are presented as mean ± SEM (n = 3-11 mice per condition). Statistical significance was determined using: (B/C, AUC) repeated-measures two-way ANOVA followed by Šidák’s multiple comparisons test (B/C % Relative change Ucn3) unpaired two-tailed Welch’s t-test. \**P* < 0.05, \*\*\**P* < 0.001, \*\*\*\**P* < 0.0001.

## Discussion

In this study, we show that when both Casp1 and Ripk3 are simultaneously inhibited in lean mice, there is an increase in insulin secretion, which results in improved glucose tolerance. These observations were made in male and female mice, using two separate genetic mouse models, in addition to pharmacological tools, in both *ex vivo* and *in vivo* conditions. While initial observations led us to interrogate the Ucn3-Sst axis, exogenous delivery of these islet hormones suggested a defect in δ-cell-mediated Sst secretion. These findings point to an unexpected basal role for both Casp1 and Ripk3 signaling in the maintenance of islet homeostasis.

We hypothesized that dual disruption of Casp1 and Ripk3 mediated an altered endocrine loop to allow for increased insulin secretion. While we determined that exogenous Ucn3 was not sufficient to hinder glucose responses in the Casp1/Ripk3 DKO or inhibitor-treated mice, the introduction of exogenous Sst (Octreotide) was equally effective in all genotypes and conditions. Unexpectedly, we measured no differences in glucose-stimulated Sst release from isolated Casp1/Ripk3 DKO islets in comparison to WT. This posed a conundrum, as we expected Sst levels to be hindered in Casp1/Ripk3 DKO mice. Importantly, we chose only representative low- and high-glucose time point samples to measure Sst following the perifusion experiments; therefore, we may not have captured differences across the entire experiment.

The roles of Casp1 and Ripk3 in whole body metabolism have been well studied, though always independently. Initial characterization led to the conclusions that blocking Casp1 expression or activity in the context of obesity had a protective effect on markers of glucose tolerance and insulin sensitivity, as well as important roles in adipogenesis [13; 42]. Conversely, other reports using similar and different Casp1 genetic models have suggested a neutral or opposite effect on metabolic outcomes [16; 30; 43]. Moreover, given the central role of Casp1 in mediating IL-1β and IL18 inflammatory responses, many reports have framed findings along this context. The importance of Ripk3 in chronic metabolic dysfunction and cardiometabolic disease is clear; deletion or inhibition of Ripk3 signaling leads to improvement in metabolic parameters [25; 28; 44; 45]. Specifically in islets and in the context of insulin release, Ripk3 activity was shown to be increased as islets underwent metabolic stress and mediated glucose intolerance in the presence of islet amyloid [25; 46]. Our initial objectives were focused on understanding the roles of Casp1 and Ripk3 individually in obesity-induced metabolic dysfunction, while also considering that inhibiting both in this setting could be protective.

When fed a 60% HFD, male but not female, single Casp1/11-deficient and Ripk3-deficient mice experienced increased adiposity that fed-forward to negatively impact glucose tolerance and insulin sensitivity, consistent with some Casp1 and all Ripk3 accounts [47]. Contrary to our initial thoughts, dual disruption of Casp1/11 and Ripk3 did not synergistically or additively affect markers of adiposity and whole-body energy metabolism. Moreover, the overall effects on glucose tolerance and insulin sensitivity were congruent with the levels of obesity as we dosed the mice based on total and not lean mass. When considering the effects on insulin secretion and glucose handling that were observed in lean, LFD-fed Casp1/11/Ripk3 DKO mice, most of these differences were not evident in the HFD-fed animals. However, in males, the glucose excursion curve of the Casp1/11/Ripk3 DKO compared to the single KO groups was observably, though not statistically, lower (Fig. 1D). In females, Casp1/11/Ripk3 DKO islets had a trending and significant increase in first and second phase insulin release in response to high glucose, respectively (Fig. S8). This could suggest that the mechanisms at play in the lean, low-fat condition that mediate higher insulin secretion are partially conserved but are mainly overwhelmed by the HFD-induced adaptations.

That insulin secretion was increased only when both Casp1 and Ripk3 were disrupted was unexpected. As stated earlier, our initial hypothesis was that Casp1/11/Ripk3 DKO mice would be protected from metabolic dysfunction due to their well-characterized roles in metabolic inflammation [9; 10]. However, while a decreased glucose excursion curve is typically suggestive of an increase in glucose tolerance, our results point to increased insulin release that is likely due to defects in endogenous Sst-mediated feedback that fail to constrain insulin secretion properly. Deletion or pharmacological inhibition of Casp1 and Ripk3 results in a maladaptive response that mirrors a hypersecretory phenotype that normally accompanies insulin resistance along the progression to T2D [48; 49]. It would be interesting to assess the phenotype of Casp1/Ripk3 DKO mice across an ageing spectrum. We would speculate that the natural age-associated insulin resistance coupled with concomitant reduced inhibition of insulin secretion pathways may predispose mice deficient for Casp1 and Ripk3 to higher levels of β cell dysfunction [50].

The canonical roles of Casp1/11 and Ripk3 are classically studied in respect to inflammation and cell death. Our initial observations of higher insulin secretion in Casp1/11/Ripk3 DKO mice begged the question as to the contributions of each of these mediators and whether their canonical functions may be responsible. While a baseline and continuous level of inflammatory signaling is critical for maintaining cellular homeostasis, the canonical roles of Casp1/11 and Ripk3 are typically engaged in response to a significant inflammatory threat [51–53]. Therefore, it remains difficult to reconcile the mechanistic link between deletion/inhibition of these proteins and the observed effects on islet biology. We were able to rule out the involvement of Casp11, important for induction of pyroptotic cell death in certain contexts by using a single Casp1 KO model that replicated the glucose excursion curves from the Casp1/11 and Ripk3 DKO. Both Casp1 and Ripk3, as a protease and kinase, respectively, have been implicated in the regulation of multiple metabolic protein targets [29; 54–59]. It would be interesting to probe whether endogenous signaling, cleavage or scaffolding roles of Casp1 and Ripk3 mediate the increased insulin secretion phenotype and the extent to which these effects may be due to alterations in various islet-resident cells.

We acknowledge the limitations and important considerations with our current study. All genetic models used are whole-body knockout mice where either Casp1/11, Casp1 or Ripk3 were deleted throughout development. It is quite possible that important compensatory changes in islet-resident or other cell types have unanticipated paracrine effects that contribute to the insulin secretion phenotype. It would be interesting, though extremely laborious, to develop mice that are specifically deficient for these factors, either in isolation or in combination in a spatial and temporal manner (i.e., β or δ cell-specific/inducible knockout models). We also acknowledge that while pharmacological inhibition supports a role for Casp1 and Ripk3 signalling in modulating insulin secretion, both *in vivo* and in isolated primary islets, there is always the potential for off target effects. However, given the effect of the inhibitors was consistent with the genetic models and did not overtly affect insulin sensitivity, we feel this supports the notion that long-term adaptation to the deletion of these genes is not a primary driver of the insulin secretion phenotype.

A final and important consideration is that the basal expression of Casp1 and Ripk3, in lean pancreatic islets, independent of overt inflammation is likely very low, with the highest expression observed in β cells [60]. This work highlights that endogenous and baseline Casp1 and Ripk3 signals act together to constrain insulin release by mediating proper endocrine signaling; however, we cannot comment as to cell-specific mechanistic contributions. Whether combined Casp1 and Ripk3 deficiency alters δ cell physiology or has pleiotropic effects on multiple islet-resident cells is something that warrants further investigation.

In conclusion, we demonstrate that in response to nutrient overload, deletion of both Casp1 and Ripk3 does not significantly protect against diet-induced obesity or metabolic dysregulation. However, in the absence of metabolic dysfunction or obesity-induced cellular stress, and independent of biological sex, Casp1 and Ripk3 signaling acts in concert to facilitate normal glucose-stimulated insulin release by maintaining sufficient Sst secretion in mice. This work highlights novel and unexpected roles for proteins that are normally assigned inflammatory relevance.

## Acknowledgements

The authors would like to thank the Animal Care and Veterinary Services facility at the University of Ottawa, as well as the University of Ottawa Louise Pelletier Histology Core Facility (RRID:SCR_021737) for the tissue embedding. Thank you to Dr. Mark Huising for helpful insight on Ucn3 biology and Dr. Vincent Poitout for help with Sst-related assays. The authors would like to apologize to colleagues whose significant work could not be included due to length, citation limitations, or author oversight.

## Funding and additional information

This work was made possible by funding provided by Diabetes Canada (OG-3-22-5657-MF) awarded to M.D.F. Durin the course of the work, M.D.G. was supported by a Canadian Institutes of Health Research (CIHR) Canadian Graduate Scholarship-Masters, C.L. was supported by a CIHR Canadian Graduate Scholarship-Masters and Canadian Islet Research Training Network NSERC CREATE, A.C. was supported by a CIHR Masters Award and a uOttawa Centre for Infection Immunity and Inflammation (CI3) Scholarship, T.K.T.S. was supported by a CIHR Vanier Scholarship, L.G. was supported by a CIHR Canadian Graduate Scholarship-Doctoral and M.H was supported by a uOttawa CI3 Scholarship. A.R.P. was supported by funding from the Canada Research Chairs Program. J.E.B. was supported by a Dorothy Killam Fellowship and the Canada Research Chairs Program.

## Supplemental Figures

**Fig. S1.**
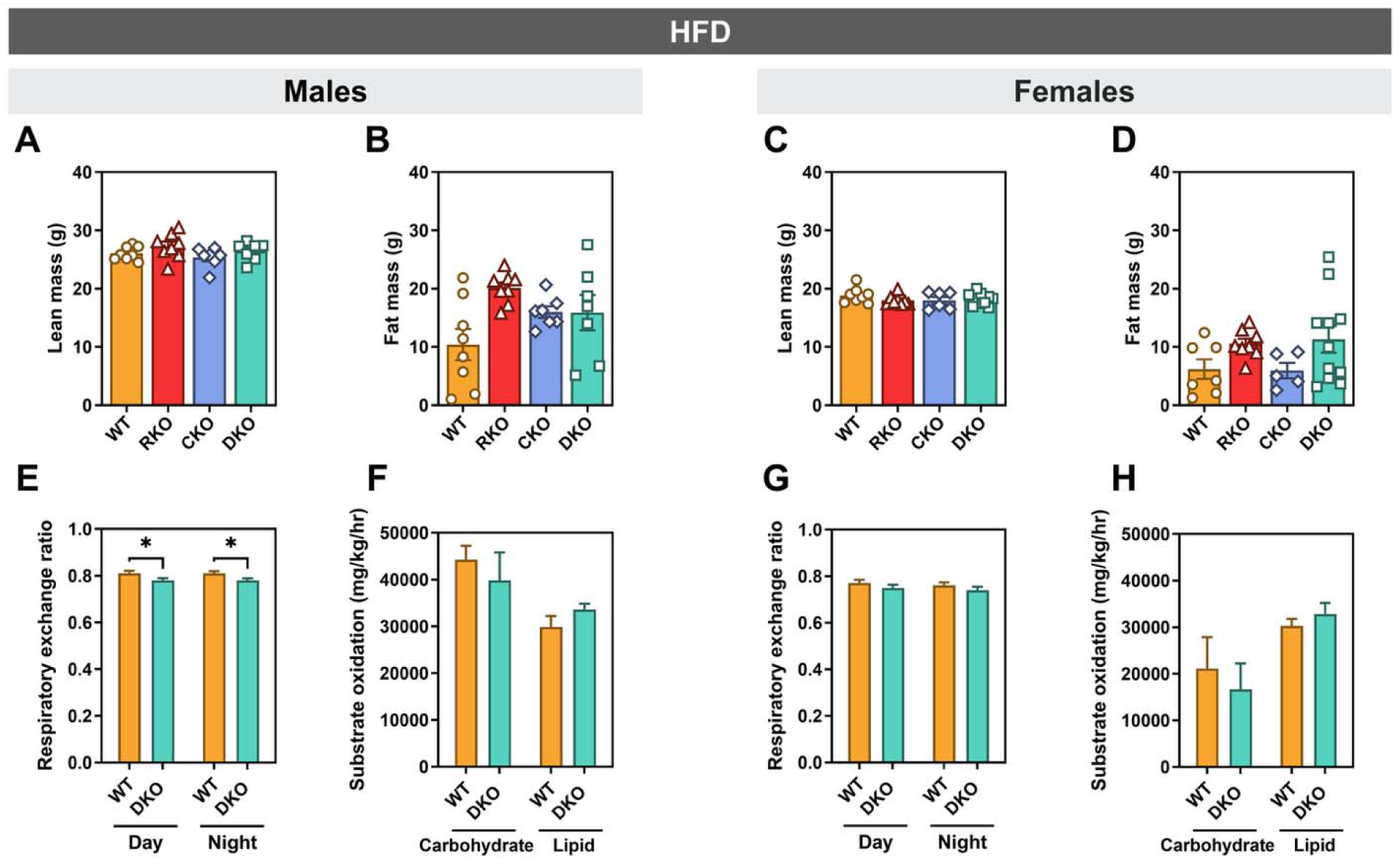
Body composition and whole-body energy metabolism are primarily unchanged between genotypes in HFD-fed male and female mice. (A-D) Lean mass and fat mass were assessed by EchoMRI-700 (EchoMRI) in WT, Ripk3 KO (RKO), Casp1/11 KO (CKO), and Casp1/11/Ripk3 DKO (DKO) mice. (E-H) Whole-body energy metabolism was assessed by indirect calorimetry using a Comprehensive Laboratory Animal Monitoring System (CLAMS; Columbus Instruments) over a 72-h period. Respiratory exchange ratio was measured during the light (day) and dark (night) cycles for WT and DKO mice (E, G). Estimated carbohydrate and lipid oxidation rates were calculated from indirect calorimetry measurements (F, H). Data are mean ± SEM (n = 4-11 mice per genotype). Statistical significance was determined by one-way ANOVA with Tukey’s multiple comparison test (A-D) and two-way ANOVA followed by Fisher’s LSD multiple comparisons test (E-H). \**P* < 0.05.

**Fig. S2.**
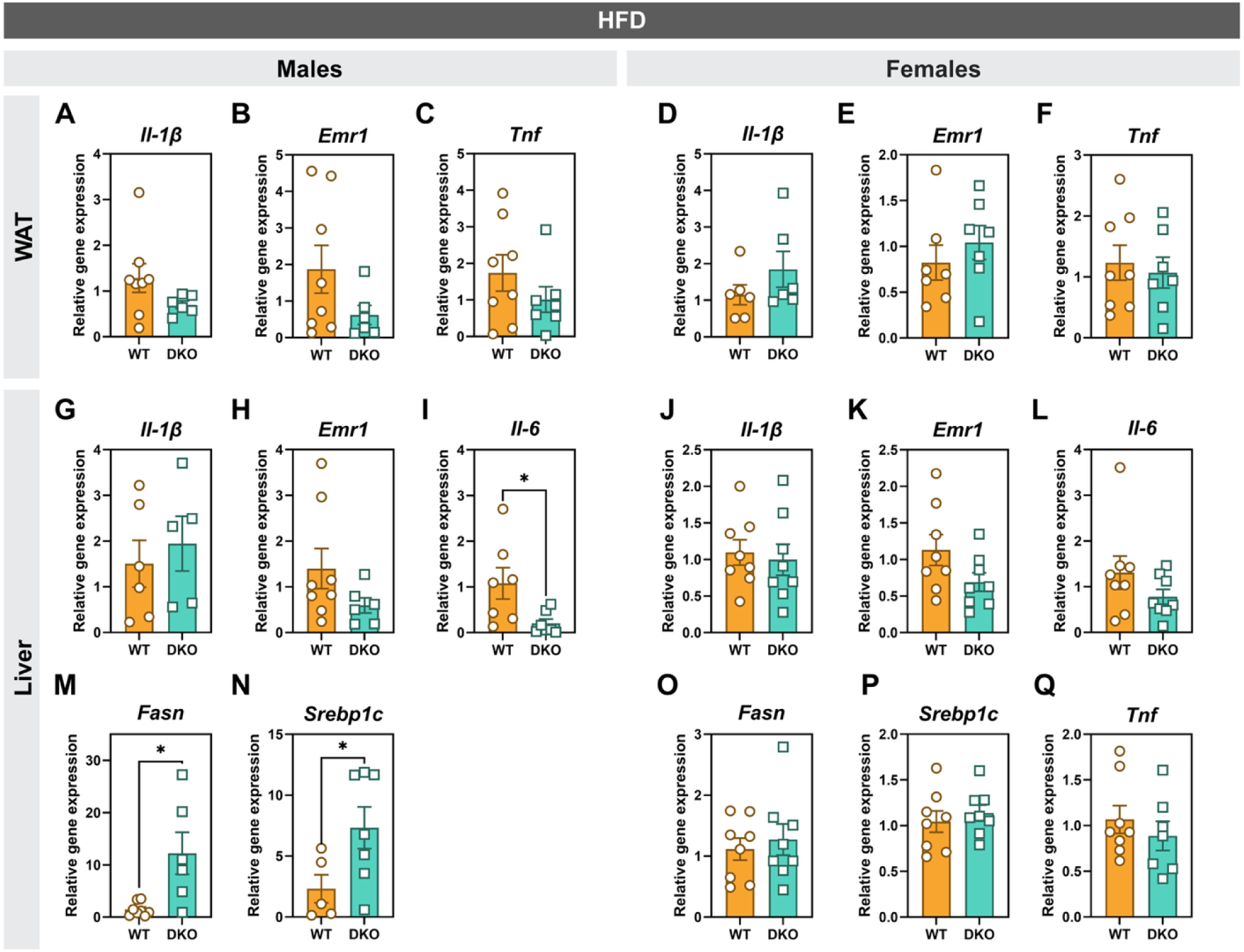
Hepatic *Il-6*, *Fasn*, and *Srebp1c* expression is altered in HFD-fed male Casp1/11/Ripk3 DKO mice, whereas WAT inflammatory gene expression is unchanged. (A-F) Relative gene expression of inflammatory transcripts (*Il-1*β, *Emr1*, and *Tnf*) in white adipose tissue (WAT) was quantified by RT-qPCR in male and female HFD-fed mice. (G-Q) Relative mRNA expression of inflammatory transcripts (*Il-1*β, *Emr1*, *Il-6*, and *Tnf* (female mice only)) and lipogenic genes (*Fasn* and *Srebp1c*) in the liver was quantified by RT-qPCR in male and female HFD-fed mice. Data are presented as mean ± SEM (n = 5-9 mice per genotype). Statistical significance was determined by unpaired two-tailed Welch’s t-test. \**P* < 0.05.

**Fig. S3.**
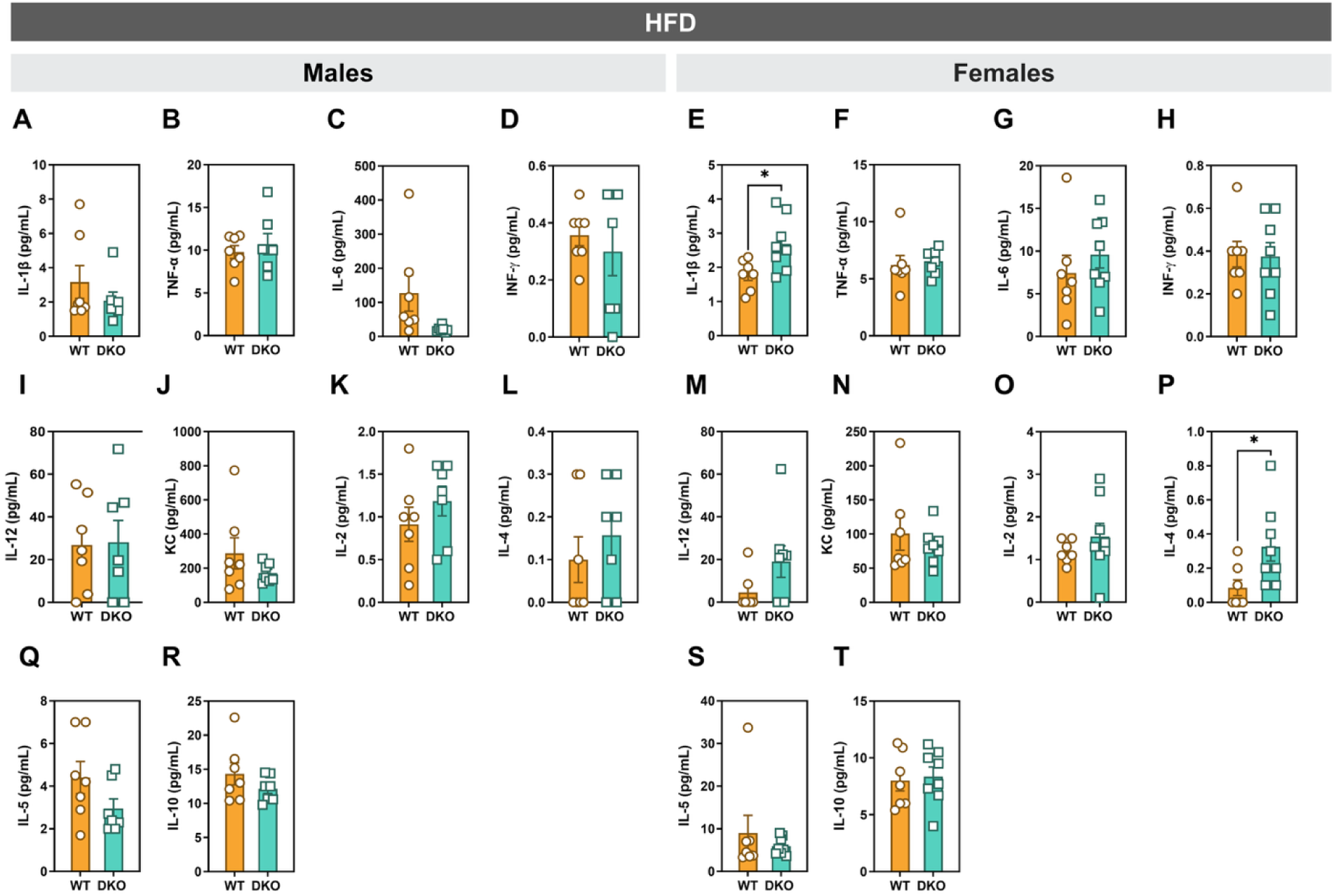
Casp1/11/Ripk3 deletion does not broadly alter circulating cytokine levels following HFD feeding. (A-T) Serum cytokine concentrations were measured by ELISA in male and female WT and Casp1/11/Ripk3 DKO mice following HFD feeding. Cytokines analyzed included IL-1β, TNF-α, IL-6, IFN-γ, IL-12, KC, IL-2, IL-4, IL-5, and IL-10. Data are presented as mean ± SEM (n = 7-8 mice per genotype). Statistical significance was determined by unpaired two-tailed Welch’s t-test. \**P* < 0.05.

**Fig. S4.**
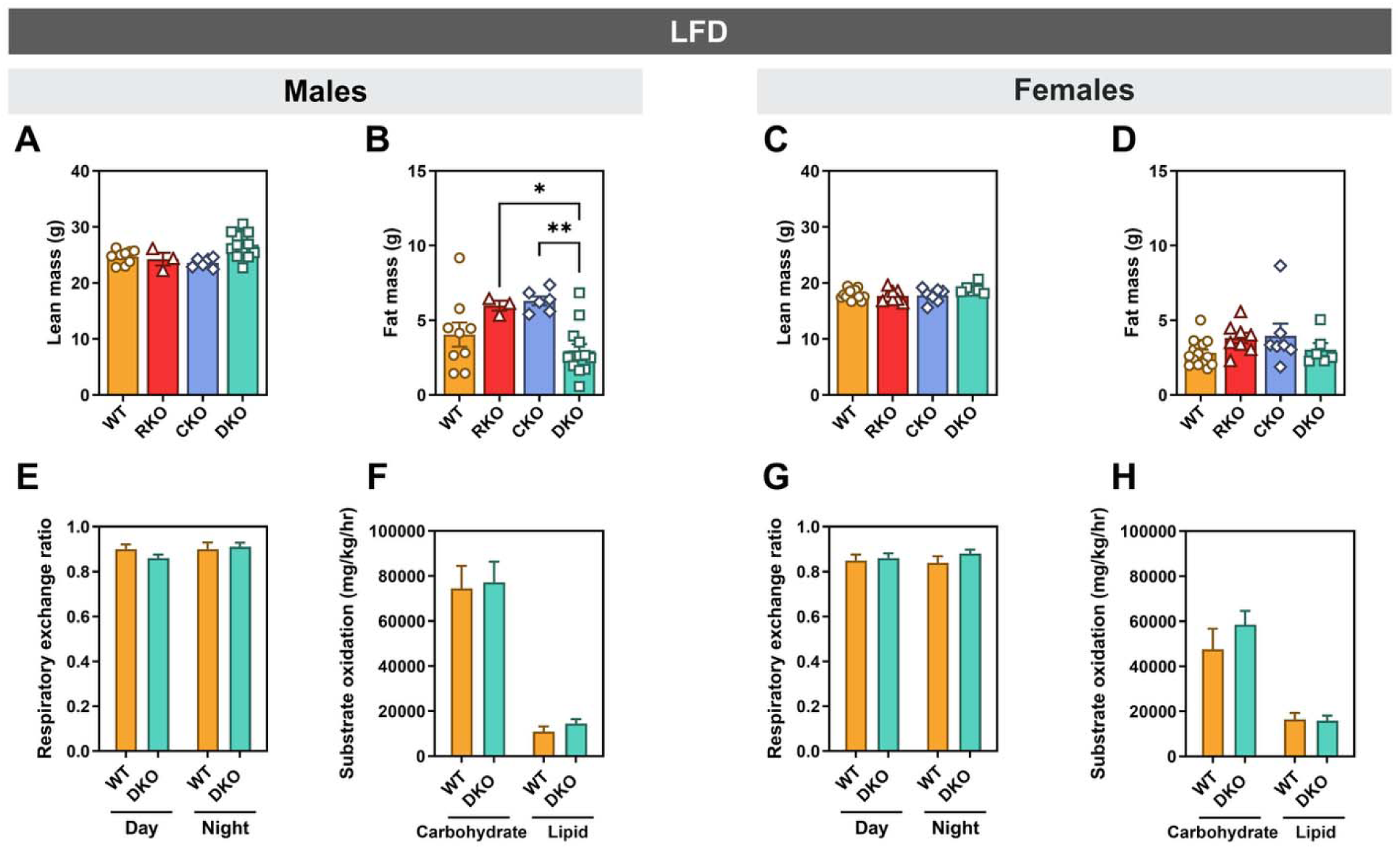
Body composition and whole-body energy metabolism are unchanged between WT and DKO in LFD-fed male and female mice. (A-D) Lean mass and fat mass were assessed by EchoMRI-700 (EchoMRI) in WT, Ripk3 KO (RKO), Casp1/11 KO (CKO), and Casp1/11/Ripk3 DKO (DKO) mice. (E-H) Whole-body energy metabolism was assessed by indirect calorimetry using a Comprehensive Laboratory Animal Monitoring System (CLAMS; Columbus Instruments) over a 72-h period. Respiratory exchange ratio was measured during the light (day) and dark (night) cycles for WT and DKO mice (E, G). Estimated carbohydrate and lipid oxidation rates were calculated from indirect calorimetry measurements (F, H). Data represents mean ± SEM (n = 3-15 mice per genotype). Statistical significance was determined by one-way ANOVA with Tukey’s multiple comparison test (A-D) and two-way ANOVA followed by Fisher’s LSD multiple comparisons test (E-H). \**P* < 0.05, \*\**P* < 0.01.

**Fig. S5.**
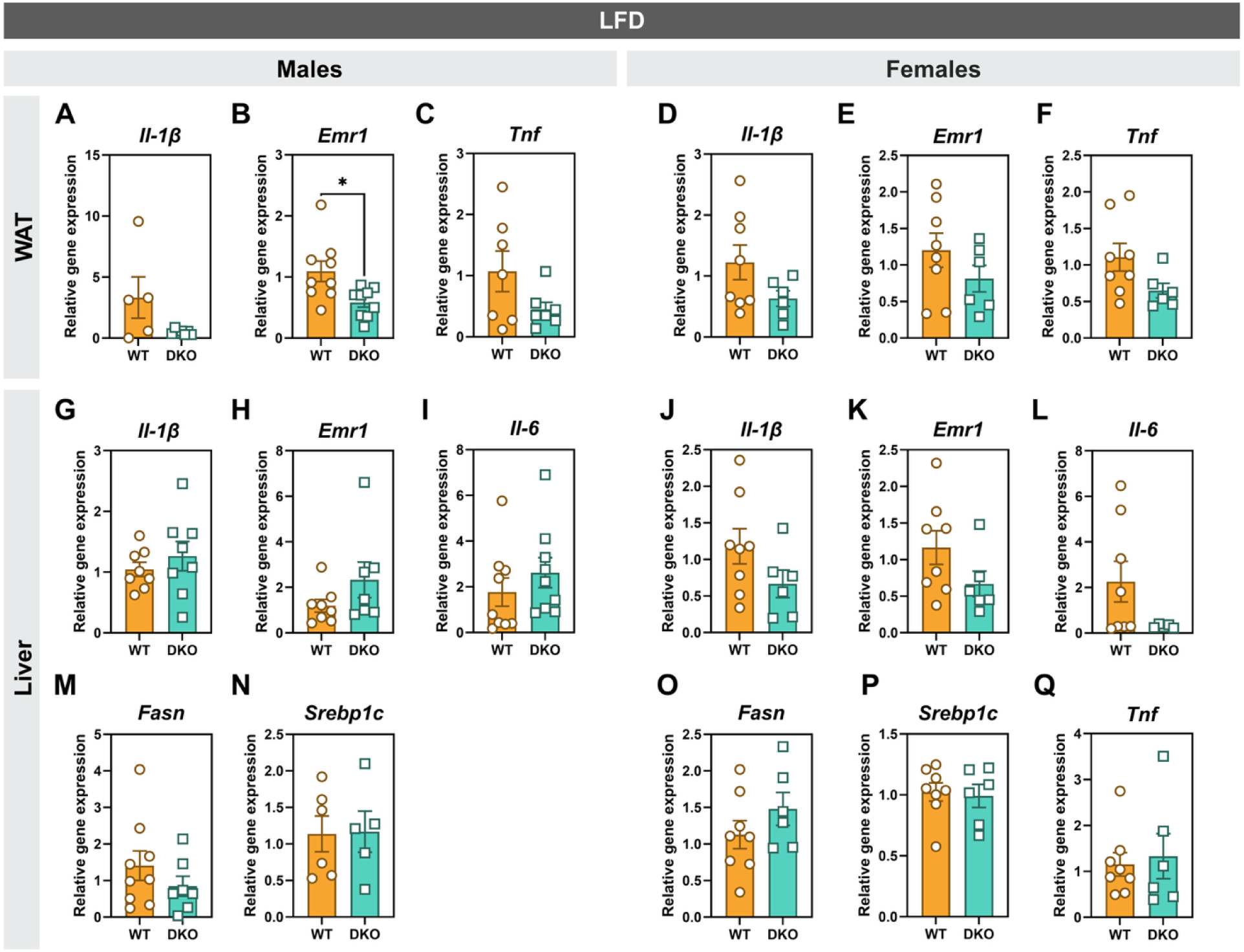
Inflammatory and lipogenic gene expression is largely preserved in the WAT and liver of LFD-fed Casp1/11/Ripk3 DKO mice. (A-F) Relative gene expression of inflammatory transcripts (*Il-1*β, *Emr1*, and *Tnf*) in white adipose tissue (WAT) was quantified by RT-qPCR in male and female LFD-fed mice. (G-Q) Relative mRNA expression of inflammatory transcripts (*Il-1*β, *Emr1*, *Il-6*, and *Tnf* (female mice only)) and lipogenic genes (*Fasn* and *Srebp1c*) in the liver was quantified by RT-qPCR in male and female LFD-fed mice. Data are presented as mean ± SEM (n = 5-9 mice per genotype). Statistical significance was determined by unpaired two-tailed Welch’s t-test. \**P* < 0.05.

**Fig. S6.**
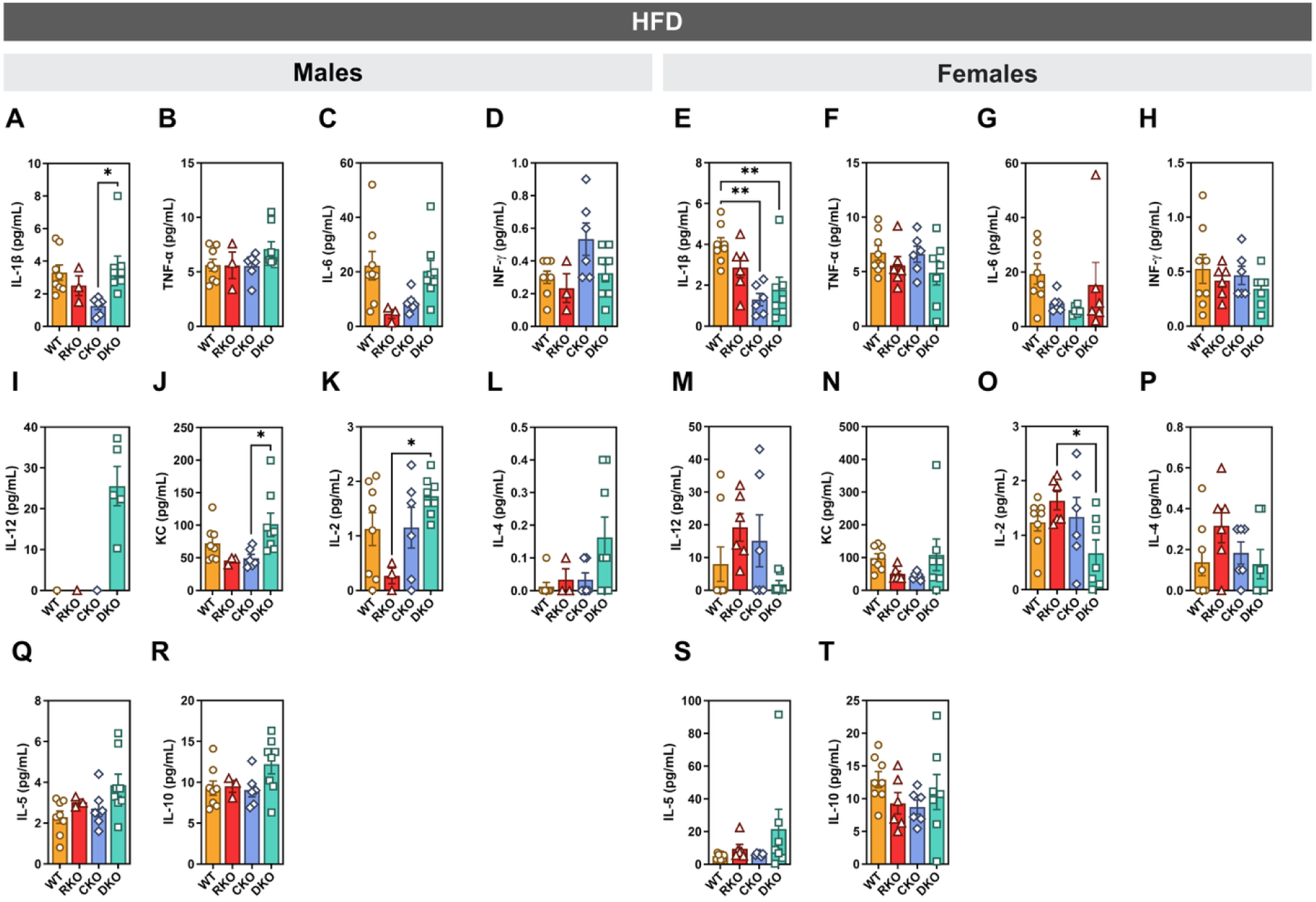
Casp1/11/Ripk3 deletion does not broadly alter circulating cytokine levels following LFD feeding. (A-T) Serum cytokine concentrations were measured in male and female WT, Ripk3 KO (RKO), Casp1/11 KO (CKO), and Casp1/11/Ripk3 DKO (DKO) mice following LFD feeding. Cytokines analyzed included IL-1β, TNF-α, IL-6, IFN-γ, IL-12, KC, IL-2, IL-4, IL-5, and IL-10. Data are presented as mean ± SEM (n = 3-8 mice per genotype). Statistical significance was determined by one-way ANOVA with Tukey’s multiple comparison test. \**P* < 0.05, \*\**P* < 0.01.

**Fig. S7.**
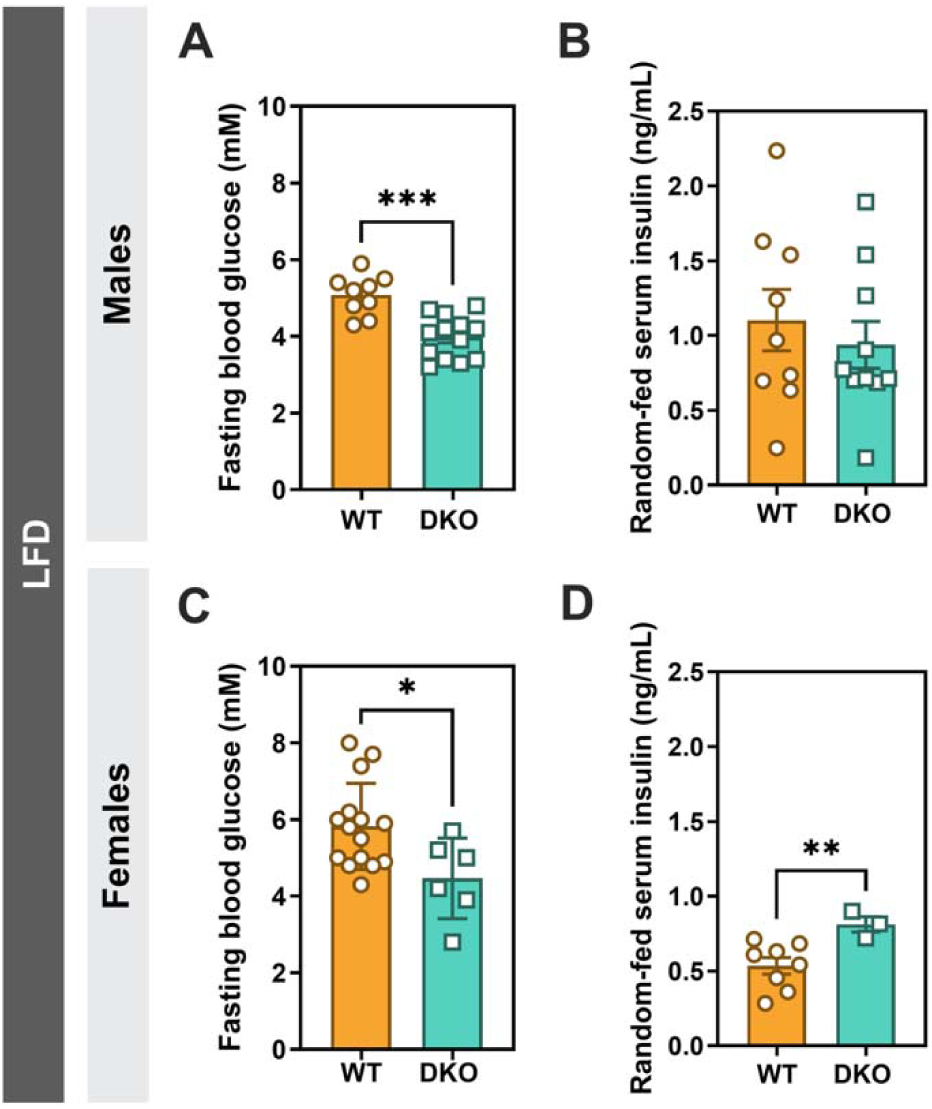
Reduced fasting serum glucose in Casp1/11/Ripk3 DKO male and female mice fed a LFD. (A, C) Serum glucose levels were measured following a ∼9 h overnight fast in WT and Casp1/11/Ripk3 DKO male and female mice fed a LFD. (B, D) Random-fed serum insulin levels were measured in WT and DKO male and female mice when given unrestricted access to food. Data are presented as mean ± SEM (n = 3-15 mice per genotype). Statistical significance was determined by unpaired two-tailed Welch’s t-test. \**P* < 0.05, \*\**P* < 0.01, \*\*\**P* < 0.001.

**Fig. S8.**
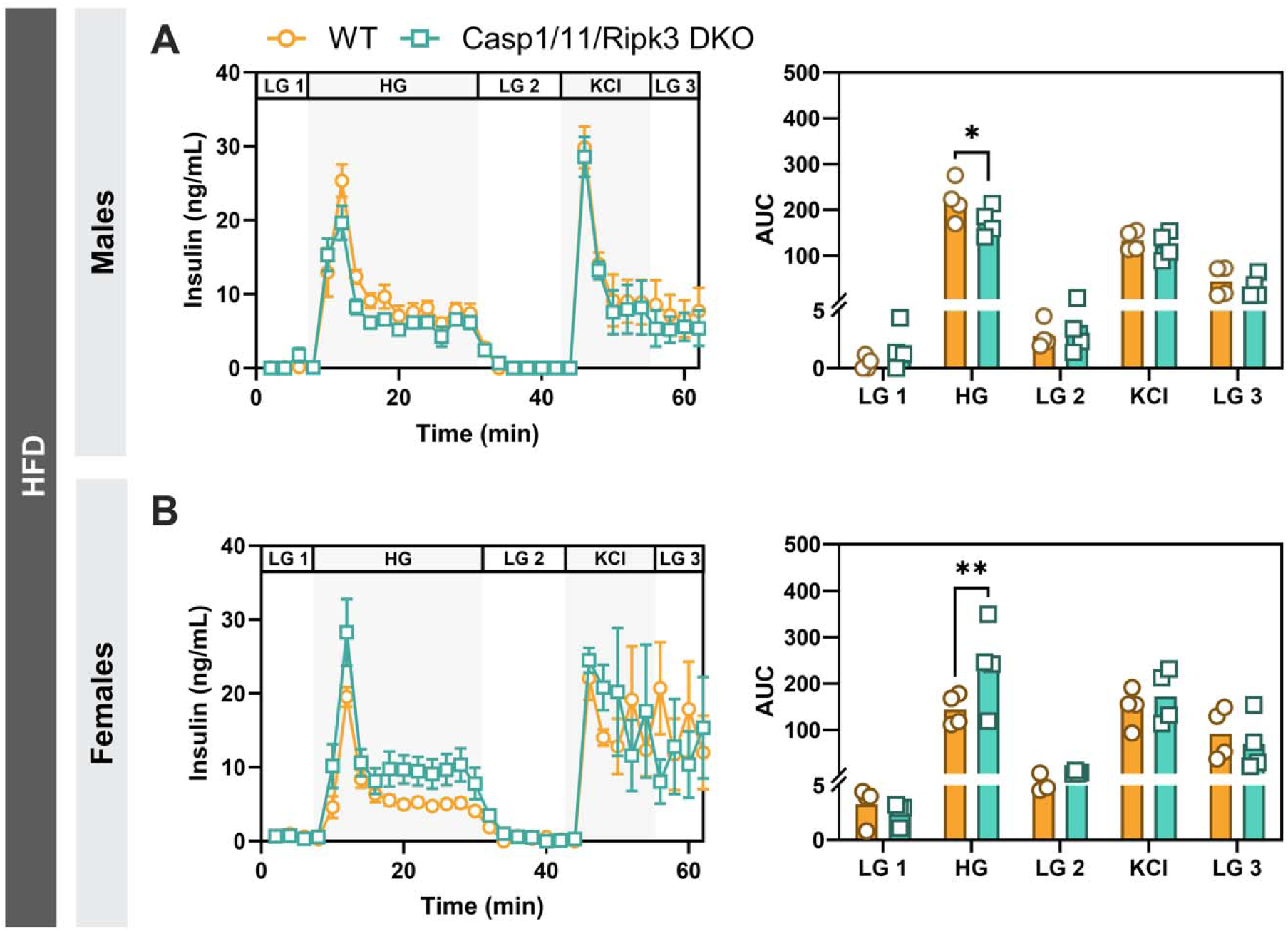
HFD feeding blunts the enhanced insulin secretion phenotype observed in LFD-fed Casp1/11/Ripk3 DKO mice. (A-B) Dynamic insulin secretion was assessed by perifusion of isolated islets from HFD-fed WT and Casp1/11/Ripk3 DKO male and female mice exposed to low glucose (LG; 2.8 mM), high glucose (HG; 16.7 mM), and depolarizing KCl (30 mM) conditions. Insulin release was measured over 62 min at a flow rate of 40 µL/min. Data are presented as mean ± SEM (n = 4 mice per genotype). Statistical significance was determined using repeated-measures two-way ANOVA followed by Šidák’s multiple comparisons test. *P < 0.05, **P < 0.01.

**Fig. S9.**
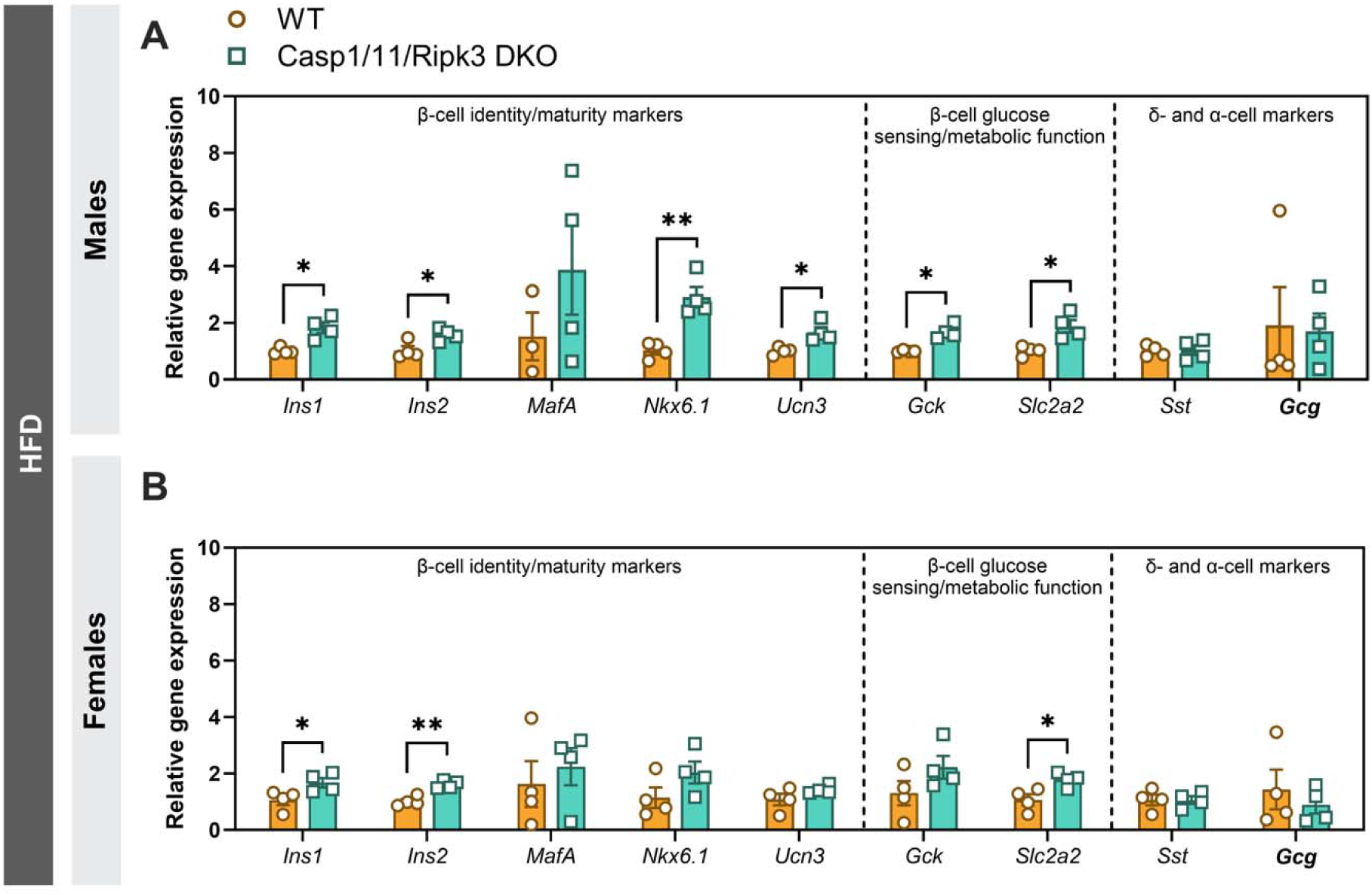
Casp1/11/Ripk3 DKO mice display altered expression of select islet transcripts following HFD feeding. (A-B) Relative mRNA expression of genes associated with β-cell identity and maturation (*Ins1*, *Ins2*, *MafA*, *Nkx6.1*, *Ucn3*), β-cell glucose sensing and metabolic function (*Gck*, *Slc2a2*), and α- and δ-cell markers (*Gcg*, *Sst*) was assessed in islets isolated from HFD-fed male and female WT and Casp1/11/Ripk3 DKO mice. Transcript abundance was quantified by RT-qPCR. Data are presented as mean ± SEM (n = 4 mice per genotype). Statistical significance was determined using an unpaired two-tailed t-test with Welch’s correction. *P < 0.05, **P < 0.01.

**Fig. S10.**
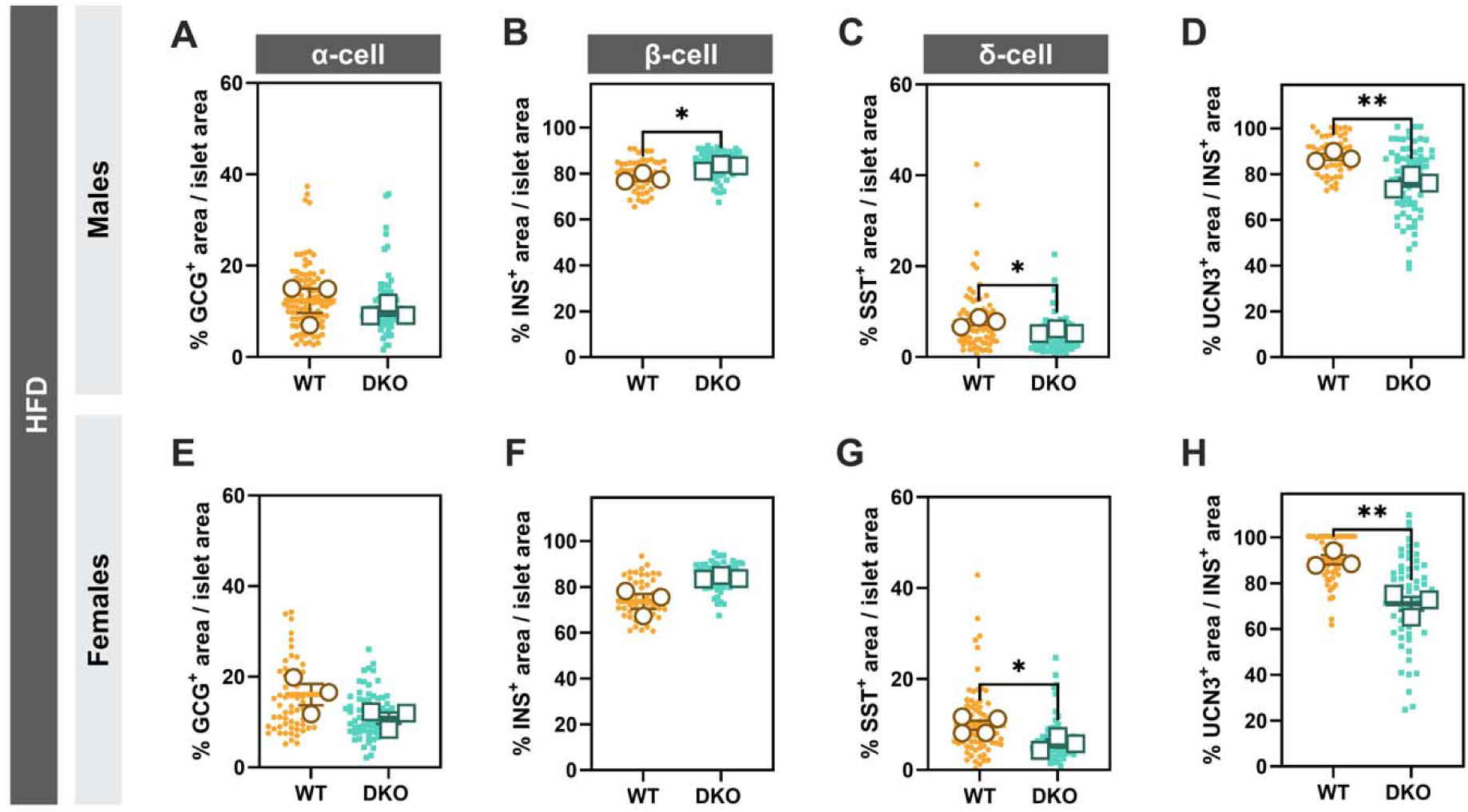
Casp1/11/Ripk3 DKO mice display changes in endocrine cell composition following HFD feeding. (A-H) Paraffin-embedded pancreatic sections from male and female mice fed a HFD were analyzed by IF staining to quantify selected islet hormones. (A, E) % glucagon (GCG) area normalized to the total area of each islet to represent the percent α-cells. (B, F) % insulin (INS) area normalized to the total area of each islet to represent the percent β-cells. Representative quantification plots from INS/urocortin 3 (UCN3) IF staining are shown. (C, G) % somatostatin (Sst) area normalized to the total area of each islet to represent the percent δ-cells. (D, H) % UCN3 area normalized to INS area per islet. Quantification of GCG, INS, and Sst area was calculated as: (hormone area / total islet area) × 100. UCN3 area was quantified as: (UCN3 area / INS area) × 100. Small dots represent individual islets (10-38 per mouse); large dots indicate the mean of all islets per mouse (n = 3 mice per genotype). Data are mean ± SEM. Statistical significance was determined by unpaired two-tailed Welch’s t-test. \**P* < 0.05, \*\**P* < 0.01.

**Fig. S11.**
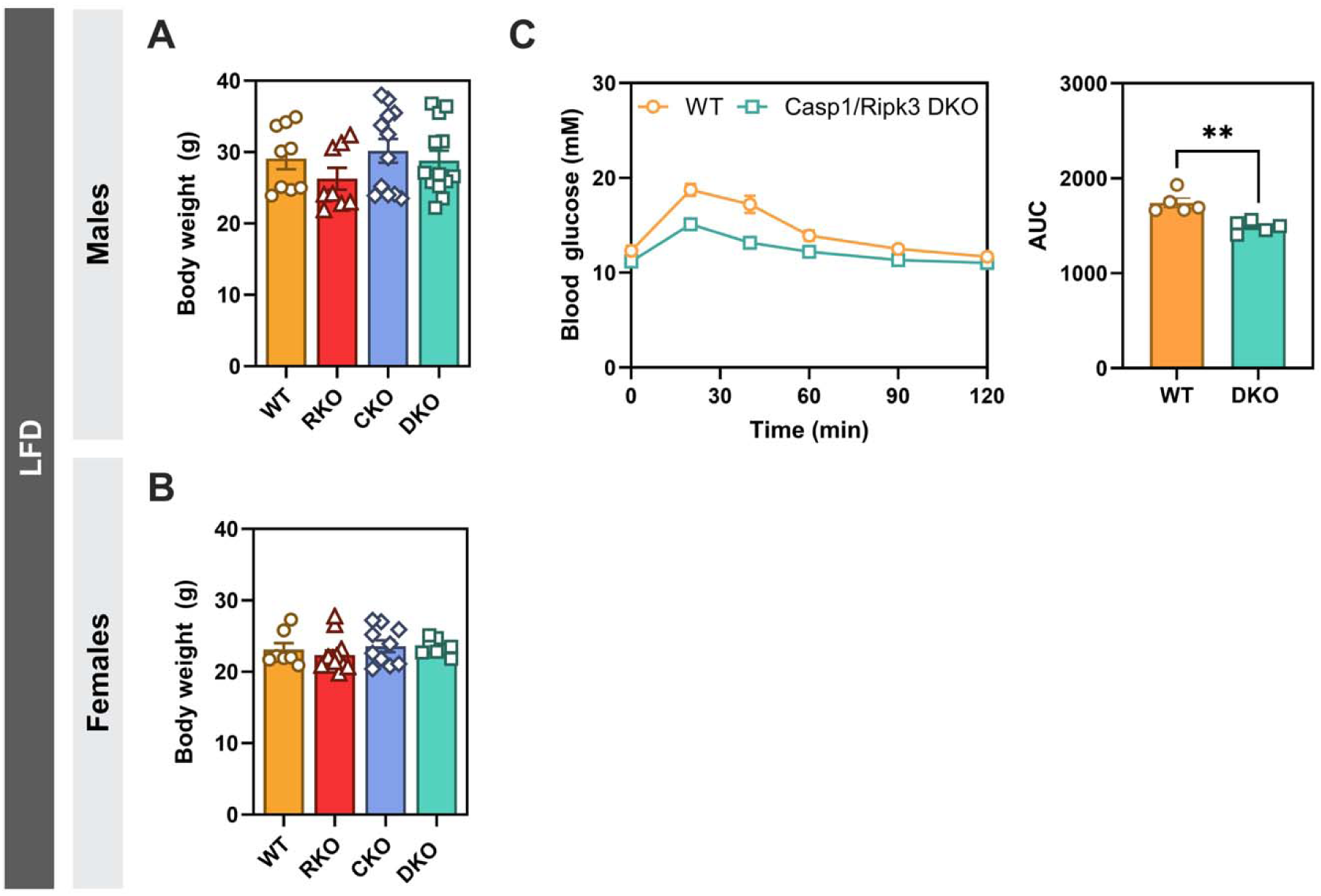
**Increased glucose tolerance is independent of Caspase 11.**(A-B) Body weight of WT, Ripk3 KO (RKO), Casp1 KO (CKO), and Casp1/Ripk3 (DKO) male and female mice fed a LFD. (C) Oral glucose tolerance test (oGTT; 2 g/kg glucose in 0.9% saline) was performed in LFD-fed mice following a 5-hour fast. Blood glucose was measured over 120 min. Data are mean ± SEM (n = 5-13 mice per genotype). Statistical significance was determined by one-way ANOVA with Tukey’s multiple comparison test (A-B) and unpaired two-tailed Welch’s t-test (C). \*\**P* < 0.01.

**Fig. S12.**
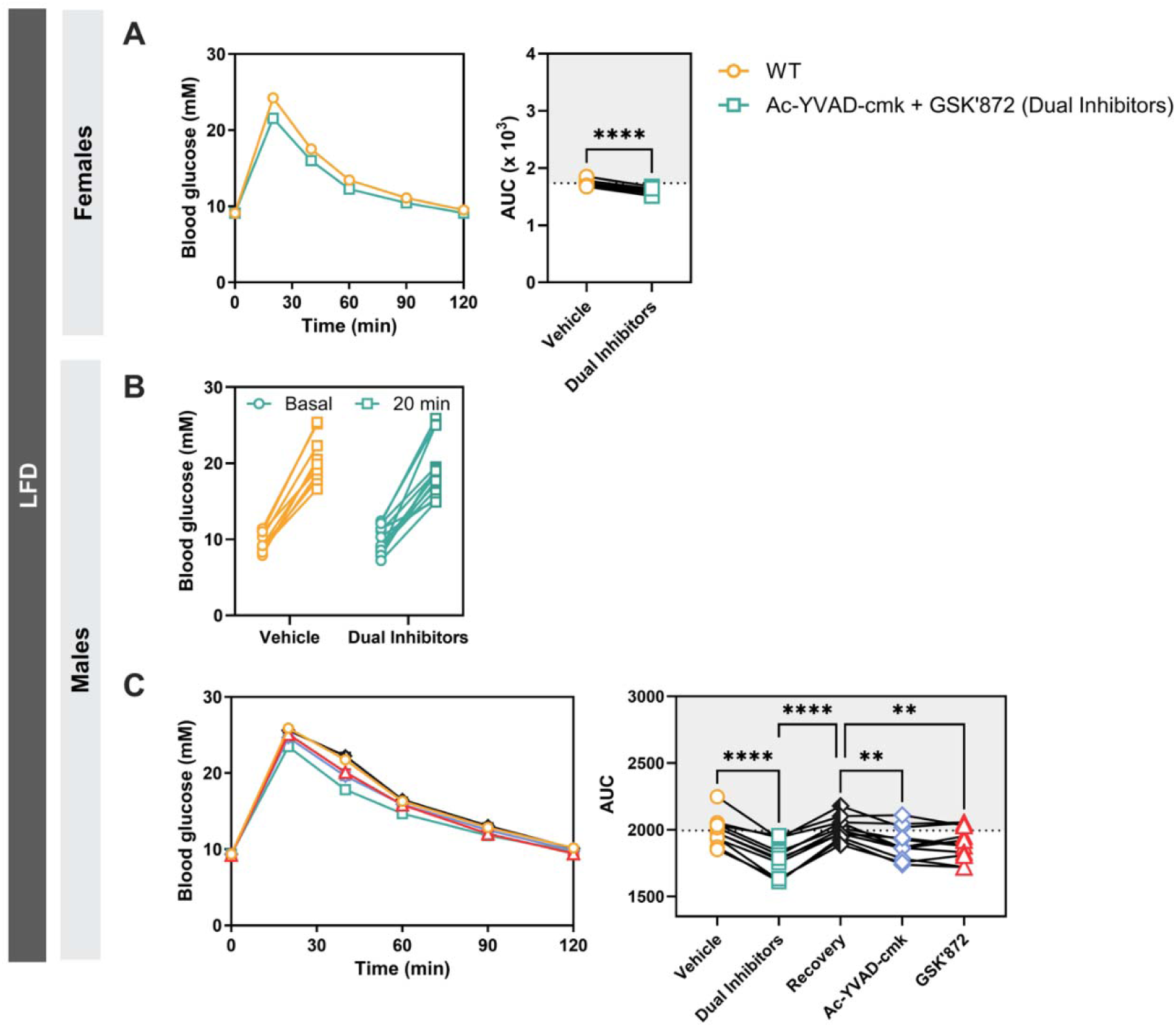
**Pharmacological inhibition of Casp1/Ripk3 recapitulates aspects of the genetic phenotype *in vivo.*** (A) Female LFD-fed C57BL/6J mice received IP injections of saline or combined Casp1/Ripk3 inhibitors (Ac-YVAD-cmk and GSK’872; 2.5 mg/kg each in 0.9% saline) over 24 h (two injections total). IP glucose tolerance tests (GTTs; 2 g/kg glucose in 0.9% saline) were performed following a 4.5-hour fast, and blood glucose levels were monitored over 120 min. (B) Male LFD-fed C57BL/6J mice received a single IP injection of saline or combined Casp1/Ripk3 inhibitors (Ac-YVAD-cmk and GSK’872; 2.5 mg/kg each in 0.9% saline) prior to IP GTTs, with blood glucose measured 20 min following glucose administration. (C) Male LFD-fed C57BL/6J mice received IP injections of saline, combined Casp1/Ripk3 inhibitors (Ac-YVAD-cmk and GSK’872; 2.5 mg/kg each in 0.9% saline), or individual Casp1 (Ac-YVAD-cmk) or Ripk3 (GSK’872) inhibitors over 24 h (two injections total). IP GTTs (2 g/kg glucose in 0.9% saline) were performed following a 4.5-hour fast, and blood glucose levels were monitored over 120 min. Data are presented as mean ± SEM (n = 9-14 mice per condition). Statistical significance was determined using a paired two-tailed t-test (A), two-way repeated-measures ANOVA followed by Fisher’s LSD multiple comparisons test (B), and repeated-measures one-way ANOVA (C). **P < 0.01, ****P < 0.0001.

**Fig. S13.**
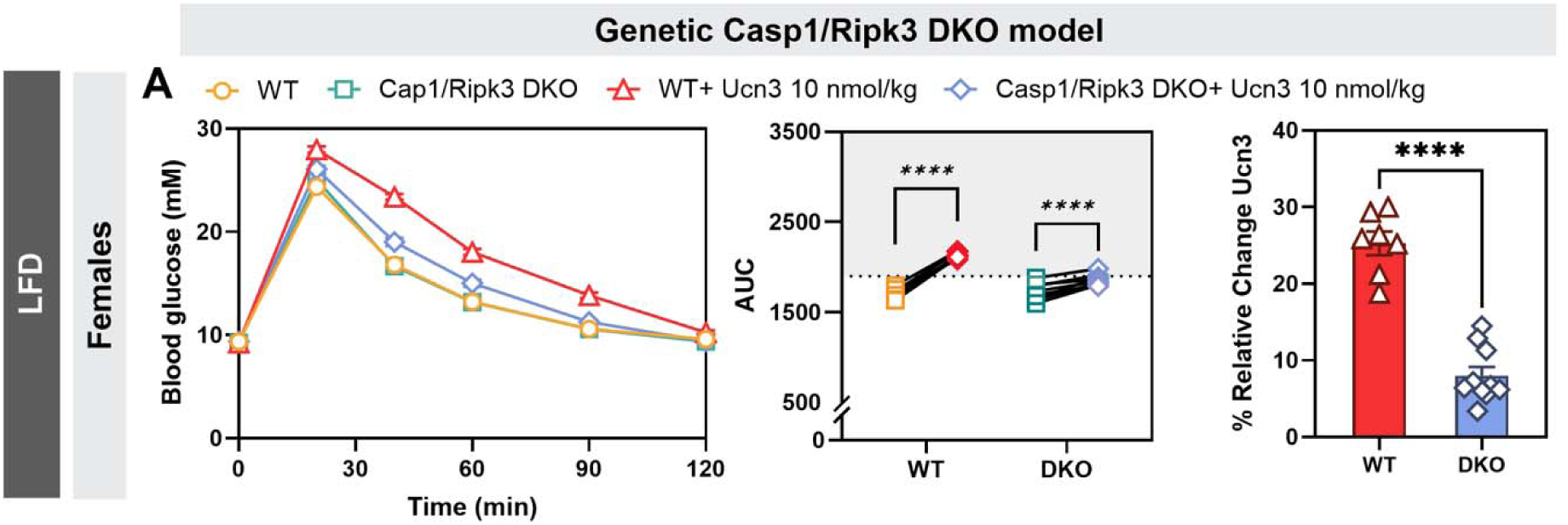
Lack of Casp1 and Ripk3 in LFD-fed female mice results in blunted Ucn3 response. (A) Following a 4.5-hour fast, female WT and Casp1/Ripk3 DKO mice were administered IP injections of saline or exogenous Ucn3 (10 nmol/kg) 5 min before an IP GTT (2 g/kg glucose in 0.9% saline). Blood glucose was monitored over 120 min. Paired AUC values were calculated for all conditions, and the percent relative change in response to UCN3 was calculated as [(UCN3 AUC - saline AUC) / saline AUC × 100]. Data are presented as mean ± SEM (n = 7-10 mice per condition). Statistical significance was determined using repeated-measures two-way ANOVA followed by Šidák’s multiple comparisons test.

**Fig. S14.**
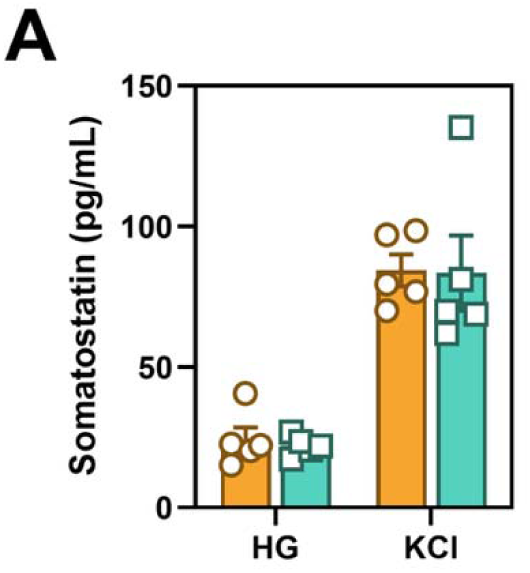
Sst secretion is unchanged in Casp1/Ripk3 DKO islets following glucose stimulation. (A) Sst secretion was measured during perifusion of isolated WT and Casp1/Ripk3 DKO islets under high glucose (HG) and KCl stimulation conditions. Data are presented as mean ± SEM (n = 5 mice per condition). Statistical significance was determined using two-way ANOVA followed by Šidák’s multiple comparisons test.

## Notes

### Competing Interest Statement

The authors have declared no competing interest.

## References

[1] Chatterjee, S., Khunti, K., Davies, M.J., 2017. Type 2 diabetes. Lancet 389(10085):2239–2251.

[2] Rawshani, A., Rawshani, A., Gudbjornsdottir, S., 2017. Mortality and Cardiovascular Disease in Type 1 and Type 2 Diabetes. N Engl J Med 377(3):300–301.

[3] Shah, A.D., Langenberg, C., Rapsomaniki, E., Denaxas, S., Pujades-Rodriguez, M., Gale, C.P., et al., 2015. Type 2 diabetes and incidence of cardiovascular diseases: a cohort study in 1.9 million people. Lancet Diabetes Endocrinol 3(2):105–113.

[4] Boni-Schnetzler, M., Mereau, H., Rachid, L., Wiedemann, S.J., Schulze, F., Trimigliozzi, K., et al., 2021. IL-1beta promotes the age-associated decline of beta cell function. iScience 24(11):103250.

[5] Ehses, J.A., Boni-Schnetzler, M., Faulenbach, M., Donath, M.Y., 2008. Macrophages, cytokines and beta-cell death in Type 2 diabetes. Biochem Soc Trans 36(Pt 3):340–342.

[6] Aguayo-Mazzucato, C., Andle, J., Lee, T.B., Jr., Midha, A., Talemal, L., Chipashvili, V., et al., 2019. Acceleration of beta Cell Aging Determines Diabetes and Senolysis Improves Disease Outcomes. Cell Metab 30(1):129–142 e124.

[7] Rojas, J., Bermudez, V., Palmar, J., Martinez, M.S., Olivar, L.C., Nava, M., et al., 2018. Pancreatic Beta Cell Death: Novel Potential Mechanisms in Diabetes Therapy. J Diabetes Res 2018:9601801.

[8] Ward, M.G., Li, G., Hao, M., 2018. Apoptotic beta-cells induce macrophage reprogramming under diabetic conditions. J Biol Chem 293(42):16160–16173.

[9] Hotamisligil, G.S., Davis, R.J., 2016. Cell Signaling and Stress Responses. Cold Spring Harb Perspect Biol 8(10).

[10] Tsalikis, J., Croitoru, D.O., Philpott, D.J., Girardin, S.E., 2013. Nutrient sensing and metabolic stress pathways in innate immunity. Cell Microbiol 15(10):1632–1641.

[11] Yuan, J., Ofengeim, D., 2024. A guide to cell death pathways. Nat Rev Mol Cell Biol 25(5):379–395.

[12] Shahzad, K., Bock, F., Al-Dabet, M.M., Gadi, I., Kohli, S., Nazir, S., et al., 2016. Caspase-1, but Not Caspase-3, Promotes Diabetic Nephropathy. J Am Soc Nephrol 27(8):2270–2275.

[13] Stienstra, R., Joosten, L.A., Koenen, T., van Tits, B., van Diepen, J.A., van den Berg, S.A., et al., 2010. The inflammasome-mediated caspase-1 activation controls adipocyte differentiation and insulin sensitivity. Cell Metab 12(6):593–605.

[14] Hotamisligil, G.S., 2006. Inflammation and metabolic disorders. Nature 444(7121):860–867.

[15] Kotas, M.E., Jurczak, M.J., Annicelli, C., Gillum, M.P., Cline, G.W., Shulman, G.I., et al., 2013. Role of caspase-1 in regulation of triglyceride metabolism. Proc Natl Acad Sci U S A 110(12):4810–4815.

[16] Kimura, H., Karasawa, T., Usui, F., Kawashima, A., Endo, Y., Kobayashi, M., et al., 2016. Caspase-1 deficiency promotes high-fat diet-induced adipose tissue inflammation and the development of obesity. Am J Physiol Endocrinol Metab 311(5):E881–E890.

[17] Park, Y.J., Warnock, G.L., Ao, Z., Safikhan, N., Meloche, M., Asadi, A., et al., 2017. Dual role of interleukin-1beta in islet amyloid formation and its beta-cell toxicity: Implications for type 2 diabetes and islet transplantation. Diabetes Obes Metab 19(5):682–694.

[18] Westwell-Roper, C.Y., Ehses, J.A., Verchere, C.B., 2014. Resident macrophages mediate islet amyloid polypeptide-induced islet IL-1beta production and beta-cell dysfunction. Diabetes 63(5):1698–1711.

[19] Westwell-Roper, C.Y., Chehroudi, C.A., Denroche, H.C., Courtade, J.A., Ehses, J.A., Verchere, C.B., 2015. IL-1 mediates amyloid-associated islet dysfunction and inflammation in human islet amyloid polypeptide transgenic mice. Diabetologia 58(3):575–585.

[20] Li, J., Xu, J., Qin, X., Yang, H., Han, J., Jia, Y., et al., 2019. Acute pancreatic beta cell apoptosis by IL-1beta is responsible for postburn hyperglycemia: Evidence from humans and mice. Biochim Biophys Acta Mol Basis Dis 1865(2):275–284.

[21] Drummer, C.t., Saaoud, F., Jhala, N.C., Cueto, R., Sun, Y., Xu, K., et al., 2023. Caspase-11 promotes high-fat diet-induced NAFLD by increasing glycolysis, OXPHOS, and pyroptosis in macrophages. Front Immunol 14:1113883.

[22] Skeldon, A.M., Morizot, A., Douglas, T., Santoro, N., Kursawe, R., Kozlitina, J., et al., 2016. Caspase-12, but Not Caspase-11, Inhibits Obesity and Insulin Resistance. J Immunol 196(1):437–447.

[23] Wada, N., Yamada, H., Motoyama, S., Saburi, M., Sugimoto, T., Kubota, H., et al., 2020. Maternal high-fat diet exaggerates diet-induced insulin resistance in adult offspring by enhancing inflammasome activation through noncanonical pathway of caspase-11. Mol Metab 37:100988.

[24] Afonso, M.B., Rodrigues, P.M., Mateus-Pinheiro, M., Simao, A.L., Gaspar, M.M., Majdi, A., et al., 2021. RIPK3 acts as a lipid metabolism regulator contributing to inflammation and carcinogenesis in non-alcoholic fatty liver disease. Gut 70(12):2359–2372.

[25] Yang, B., Maddison, L.A., Zaborska, K.E., Dai, C., Yin, L., Tang, Z., et al., 2020. RIPK3-mediated inflammation is a conserved beta cell response to ER stress. Sci Adv 6(51).

[26] Zhou, X., Xie, L., Xia, L., Bergmann, F., Buchler, M.W., Kroemer, G., et al., 2017. RIP3 attenuates the pancreatic damage induced by deletion of ATG7. Cell Death Dis 8(7):e2918.

[27] Karunakaran, D., Geoffrion, M., Wei, L., Gan, W., Richards, L., Shangari, P., et al., 2016. Targeting macrophage necroptosis for therapeutic and diagnostic interventions in atherosclerosis. Sci Adv 2(7):e1600224.

[28] Lin, J., Li, H., Yang, M., Ren, J., Huang, Z., Han, F., et al., 2013. A role of RIP3-mediated macrophage necrosis in atherosclerosis development. Cell Rep 3(1):200–210.

[29] Shao, W., Yeretssian, G., Doiron, K., Hussain, S.N., Saleh, M., 2007. The caspase-1 digestome identifies the glycolysis pathway as a target during infection and septic shock. J Biol Chem 282(50):36321–36329.

[30] Sun, Q., Scott, M.J., 2016. Caspase-1 as a multifunctional inflammatory mediator: noncytokine maturation roles. J Leukoc Biol 100(5):961–967.

[31] Denes, A., Lopez-Castejon, G., Brough, D., 2012. Caspase-1: is IL-1 just the tip of the ICEberg? Cell Death Dis 3:e338.

[32] Islam, T., Afonso, M.B., Rodrigues, C.M.P., 2022. The role of RIPK3 in liver mitochondria bioenergetics and function. Eur J Clin Invest 52(3):e13648.

[33] Kuida, K., Lippke, J.A., Ku, G., Harding, M.W., Livingston, D.J., Su, M.S., et al., 1995. Altered cytokine export and apoptosis in mice deficient in interleukin-1 beta converting enzyme. Science 267(5206):2000–2003.

[34] Newton, K., Sun, X., Dixit, V.M., 2004. Kinase RIP3 is dispensable for normal NF-kappa Bs, signaling by the B-cell and T-cell receptors, tumor necrosis factor receptor 1, and Toll-like receptors 2 and 4. Mol Cell Biol 24(4):1464–1469.

[35] Rauch, I., Deets, K.A., Ji, D.X., von Moltke, J., Tenthorey, J.L., Lee, A.Y., et al., 2017. NAIP-NLRC4 Inflammasomes Coordinate Intestinal Epithelial Cell Expulsion with Eicosanoid and IL-18 Release via Activation of Caspase-1 and −8. Immunity 46(4):649–659.

[36] Mina, A.I., LeClair, R.A., LeClair, K.B., Cohen, D.E., Lantier, L., Banks, A.S., 2018. CalR: A Web-Based Analysis Tool for Indirect Calorimetry Experiments. Cell Metab 28(4):656–666 e651.

[37] Hoyeck, M.P., Angela Ching, M.E., Basu, L., van Allen, K., Palaniyandi, J., Perera, I., et al., 2024. The aryl hydrocarbon receptor in beta-cells mediates the effects of TCDD on glucose homeostasis in mice. Mol Metab 81:101893.

[38] Ibrahim, M., MacFarlane, E.M., Matteo, G., Hoyeck, M.P., Rick, K.R.C., Farokhi, S., et al., 2020. Functional cytochrome P450 1A enzymes are induced in mouse and human islets following pollutant exposure. Diabetologia 63(1):162–178.

[39] Livak, K.J., Schmittgen, T.D., 2001. Analysis of relative gene expression data using real-time quantitative PCR and the 2(-Delta Delta C(T)) Method. Methods 25(4):402–408.

[40] Abu Khweek, A., Amer, A.O., 2020. Pyroptotic and non-pyroptotic effector functions of caspase-11. Immunol Rev 297(1):39–52.

[41] van der Meulen, T., Donaldson, C.J., Caceres, E., Hunter, A.E., Cowing-Zitron, C., Pound, L.D., et al., 2015. Urocortin3 mediates somatostatin-dependent negative feedback control of insulin secretion. Nat Med 21(7):769–776.

[42] Morrison, M.C., Mulder, P., Salic, K., Verheij, J., Liang, W., van Duyvenvoorde, W., et al., 2016. Intervention with a caspase-1 inhibitor reduces obesity-associated hyperinsulinemia, non-alcoholic steatohepatitis and hepatic fibrosis in LDLR-/-.Leiden mice. Int J Obes (Lond) 40(9):1416–1423.

[43] Wang, H., Capell, W., Yoon, J.H., Faubel, S., Eckel, R.H., 2014. Obesity development in caspase-1-deficient mice. Int J Obes (Lond) 38(1):152–155.

[44] Gautheron, J., Vucur, M., Reisinger, F., Cardenas, D.V., Roderburg, C., Koppe, C., et al., 2014. A positive feedback loop between RIP3 and JNK controls non-alcoholic steatohepatitis. EMBO Mol Med 6(8):1062–1074.

[45] He, S., Wang, L., Miao, L., Wang, T., Du, F., Zhao, L., et al., 2009. Receptor interacting protein kinase-3 determines cellular necrotic response to TNF-alpha. Cell 137(6):1100–1111.

[46] Mukherjee, N., Contreras, C.J., Lin, L., Colglazier, K.A., Mather, E.G., Kalwat, M.A., et al., 2024. RIPK3 promotes islet amyloid-induced beta-cell loss and glucose intolerance in a humanized mouse model of type 2 diabetes. Mol Metab 80:101877.

[47] !!! INVALID CITATION !!! [16; 24-28; 43; 44; 46-48].

[48] Dankner, R., Chetrit, A., Shanik, M.H., Raz, I., Roth, J., 2009. Basal-state hyperinsulinemia in healthy normoglycemic adults is predictive of type 2 diabetes over a 24-year follow-up: a preliminary report. Diabetes Care 32(8):1464–1466.

[49] Thomas, D.D., Corkey, B.E., Istfan, N.W., Apovian, C.M., 2019. Hyperinsulinemia: An Early Indicator of Metabolic Dysfunction. J Endocr Soc 3(9):1727–1747.

[50] Koopman, R.J., Mainous, A.G., 3rd, Diaz, V.A., Geesey, M.E., 2005. Changes in age at diagnosis of type 2 diabetes mellitus in the United States, 1988 to 2000. Ann Fam Med 3(1):60–63.

[51] Ruhl, S., Broz, P., 2015. Caspase-11 activates a canonical NLRP3 inflammasome by promoting K(+) efflux. Eur J Immunol 45(10):2927–2936.

[52] Bauernfeind, F.G., Horvath, G., Stutz, A., Alnemri, E.S., MacDonald, K., Speert, D., et al., 2009. Cutting edge: NF-kappaB activating pattern recognition and cytokine receptors license NLRP3 inflammasome activation by regulating NLRP3 expression. J Immunol 183(2):787–791.

[53] Orozco, S.L., Daniels, B.P., Yatim, N., Messmer, M.N., Quarato, G., Chen-Harris, H., et al., 2019. RIPK3 Activation Leads to Cytokine Synthesis that Continues after Loss of Cell Membrane Integrity. Cell Rep 28(9):2275–2287 e2275.

[54] Kahns, S., Kalai, M., Jakobsen, L.D., Clark, B.F., Vandenabeele, P., Jensen, P.H., 2003. Caspase-1 and caspase-8 cleave and inactivate cellular parkin. J Biol Chem 278(26):23376–23380.

[55] Niu, Z., Shi, Q., Zhang, W., Shu, Y., Yang, N., Chen, B., et al., 2017. Caspase-1 cleaves PPARgamma for potentiating the pro-tumor action of TAMs. Nat Commun 8(1):766.

[56] Denes, A., Lopez-Castejon, G., Brough, D., 2012. Caspase-1: is IL-1 just the tip of the ICEberg? Cell Death Dis 3(7):e338.

[57] Miggin, S.M., Palsson-McDermott, E., Dunne, A., Jefferies, C., Pinteaux, E., Banahan, K., et al., 2007. NF-kappaB activation by the Toll-IL-1 receptor domain protein MyD88 adapter-like is regulated by caspase-1. Proc Natl Acad Sci U S A 104(9):3372–3377.

[58] Yang, Z., Wang, Y., Zhang, Y., He, X., Zhong, C.Q., Ni, H., et al., 2018. RIP3 targets pyruvate dehydrogenase complex to increase aerobic respiration in TNF-induced necroptosis. Nat Cell Biol 20(2):186–197.

[59] Wu, L., Zhang, X., Zheng, L., Zhao, H., Yan, G., Zhang, Q., et al., 2020. RIPK3 Orchestrates Fatty Acid Metabolism in Tumor-Associated Macrophages and Hepatocarcinogenesis. Cancer Immunol Res 8(5):710–721.

[60] Guo, J., Fu, W., 2020. Immune regulation of islet homeostasis and adaptation. J Mol Cell Biol 12(10):764–774.

